# Experimental Evolution of Yeast Reveals Trade-offs Between Early and Late Stationary Phase

**DOI:** 10.64898/2026.03.12.711341

**Authors:** Jason Tarkington, Aditya Mahadevan, Angelina G. Chan, Gavin Sherlock

## Abstract

Stationary phase in yeast and other microorganisms begins when a limiting nutrient in the environment is exhausted and cell division ceases. Most cells subsequently enter quiescence and lose viability. In spent media, without metabolic byproducts being diluted, cellular processes can modify the environment and cause the relative growth rates of different genotypes to vary over the course of stationary phase. In this work we experimentally evolve *S. cerevisiae* in batch culture, varying the time spent in stationary phase between growth cycles. We measure the relative fitness of the resulting adaptive clones across a range of environments: with different amounts of time in stationary phase and in two different carbon sources. By comparing the inferred performance (relative growth rate during a period of the growth cycle) of a mutant to that of its ancestor, we can estimate the effects of each observed mutation on performance during various phases of growth. We show that when an adaptive mutation emerges in growth cycles that include a stationary phase, its effect on stationary phase performance is largely independent of the type of carbon source provided. However, for the same group of mutants, mutational effects on performance in early stationary phase are negatively correlated with those effects in late stationary phase, suggesting a trade-off. We also show that increased intervals of stationary phase result in larger fitness effects of adaptive mutations and distinct routes of adaptation. Together, these results demonstrate that stationary phase consists of more than one distinct fitness-related phenotype, and that the phenotypes that allow for high performance in the first few days of stationary phase trade off with those that allow for high performance in later stationary phase.

## Introduction

The vast majority of microbes exists in a non-proliferative state often referred to as stationary phase—though there is surely great diversity within this broad classification (Lennon & Jones, 2011; Stevenson, 1977). The distribution of life-sustaining resources, particularly in the oceans, promotes a feast or famine lifestyle (Behringer et al., 2024; Hobbie & Hobbie, 2013; Reese et al., 2018; Zhang & Rubin, 2013) in which microbes must survive long periods of time without resources. Stationary phase begins after resources are depleted, triggering entry into a quiescent state (Kolter et al., 1993; Nyström, 2004; Werner-Washburne et al., 1993; Werner-Washburne et al., 1996), wherein cells turn down expression of genes involved in cell-division and turn on expression of genes involved in responding to stress and cellular maintenance (De Virgilio, 2012; Galdieri et al., 2010; Herman, 2002; Miles et al., 2021; Mitra et al., 2025). Subpopulations of cells that do not enter this quiescent state either die or continue to divide slowly by scavenging resources from their dying neighbors, which can itself give rise to a range of rich evolutionary dynamics (Finkel, 2006; Finkel & Kolter, 1999; Sun & Gresham, 2021; Zambrano et al., 1993; Zinser & Kolter, 2004). Understanding the fate of these nutrient-limited cells has important evolutionary and ecological implications, as microbes play a vital role in global carbon and nitrogen cycles (Armbrust, 2009; Azam et al., 1994; Higgins et al., 2012), shape Earth’s atmosphere and climate (Bardgett et al., 2008), drive marine food webs (Worden et al., 2015), and make up a substantial portion of global biomass (Bar-On et al., 2018).

Although microbes spend most of their time in stationary phase, growth rate and colony size are often used as a proxy for cellular “fitness” (Andersson & Hughes, 2010; Giaever et al., 2002; Maier et al., 2018; Miller et al., 2022; Nilsson et al., 2006; Paulander et al., 2007; Sandegren et al., 2008; Walkiewicz et al., 2012). Particularly in laboratory conditions (e.g. batch culture with serial dilution), exponential growth rate and time spent in lag phase have the dominant effect on a lineage’s differential growth rate: fitness gains during experimental evolution are primarily derived from improvements in these traits (Vasi et al., 1994). Because cells are either constantly fed (e.g., Dykhuizen & Hartl, 1983; Gresham & Hong, 2015) or regularly transferred to fresh medium (e.g., Johnson et al., 2021; Lenski et al., 1991), survival in stationary phase is a less important component of cellular fitness. In nature, however, where most microbes are nutrient-limited, survival during stationary phase is a vital component of cellular fitness.

Adaptation to long periods of stationary phase could function by increasing the survival time of quiescent cells and slowing chronological aging, or by increasing the replicative fitness of cells via an increased ability to metabolize nutrients found in dead cells or spent media. In *Escherichia coli*, mutants with a <u>g</u>rowth <u>a</u>dvantage in <u>s</u>tationary <u>p</u>hase (GASP) emerge during prolonged starvation and eventually take over stationary phase cultures (Zambrano et al., 1993). These GASP mutants have a growth advantage in stationary phase due to their enhanced catabolic capacity to metabolize dead cells and cellular debris (Zinser & Kolter, 1999). Observations of GASP-like phenotypes are less common in eukaryotes, but adaptive regrowth has been observed in stationary phase yeast cultures (Fabrizio et al., 2004). GASP-like phenotypes have also been observed in two yeast isolates recovered from 2-year-old sealed bottles of beer, both of which contained many mutations in the TOR complex (Aouizerat et al., 2019). Mutations in *HSP90*, an important molecular chaperone, have also been shown to slow chronological aging and increase survival in stationary phase cultures (Harris et al., 2001). Many adaptive yeast mutants with increased stationary phase performance have been isolated from populations evolved with growth cycles that included 2 days of stationary phase. For example, autodiploidization, chromosome 11 duplications, and *SXM1*, *FPK1*, and *SSK1* mutations were all positively selected in these evolutionary conditions (Y. Li et al., 2019).

Adaptations to stationary phase can also have an impact outside of stationary phase. For example, many mutants display a trade-off between growth rate and survival time (Biselli et al., 2020; De Paepe & Taddei, 2006; Goldhill & Turner, 2014; Kvitek & Sherlock, 2013; Y. Li et al., 2018; Zhu et al., 2023). Such trade-offs are crucial for determining the long-term fates of mutations, particularly in environments that change over time (Ferenci, 2016; Stearns, 1989). Survival during stationary phase is often treated as a singular trait and measured as a single metric fitted from multiple measurements during a predefined period following the cessation of growth (De Paepe & Taddei, 2006; Sun et al., 2020). However, the death rate of stationary phase cells is not constant over time (Minois et al., 2009; Vasi et al., 1994), suggesting that there may be uncharacterized trade-offs between performance in these different intervals of stationary phase. We sought to determine how time spent in stationary phase impacts evolutionary dynamics and outcomes. The harsh conditions associated with stationary phase may result in stronger selection and a faster loss of diversity (Hartl & Clark, 2007; Smith & Haigh, 1974). Alternatively, the persistence of cells on the byproducts of other cells via cross-feeding could produce ecological niches (Rozen & Lenski, 2000; Tilman, 1982) which sustain diversity (Ascanio et al., 2024; Chesson, 2011; Levine & Hille Ris Lambers, 2009; Sexton et al., 2017).

We evolved replicate populations of barcoded yeast across a series of growth cycles of variable length, thereby tuning the amount of time spent in stationary phase. We performed whole genome sequencing on hundreds of adaptive mutants isolated from these evolutions. Then we measured the fitnesses of these clones in conditions that allowed us to estimate stationary phase performance following growth on either (1) glucose or (2) Glycerol plus Ethanol (0.5% glycerol + 0.5% ethanol, hereafter referred to as Gly/Eth), in 2-day intervals following resource exhaustion in the Gly/Eth-limited medium. We found that performances in stationary phase (SP) following glucose versus Gly/Eth exhaustion were positively correlated, indicating an overlap in the underlying traits that contribute to these two performance metrics. By contrast, we found a negative correlation between early and late stationary phase performances, indicating a tradeoff. This finding challenges the monolithic view of stationary phase performance and suggests that stationary phase should be broken down temporally to fully appreciate the complexities of adaptation in resource-limited environments.

## Results

### Experimental Overview

We evolved barcoded yeast in triplicate by serial batch transfer in a non-fermentable Gly/Eth medium, varying the time between transfers and therefore the amount of time spent in stationary phase, which ranged from 0 to 8 days. The serial dilution factor was ∼1:250 and ∼5 × 10^7^ cells were transferred between cycles allowing for roughly 8 generations per cycle. Cultures grew to their maximum density in 2 days after which they entered stationary phase. We tracked the populations’ evolutionary dynamics by estimating the fractional abundance of each of the ∼500,000 barcoded lineages at regular intervals via barcode sequencing (Fig. 1 and Fig. S1). The resulting lineage trajectories were then used to select timepoints from each evolution at which there was both sufficient adaptation and diversity for isolating adaptive clones. Thousands of clones were isolated across all evolutionary conditions and their barcodes were sequenced. A set of 480 uniquely barcoded clones that were likely to be adaptive (based on their barcode trajectories) was identified and their genomes sequenced.

**Figure 1.**
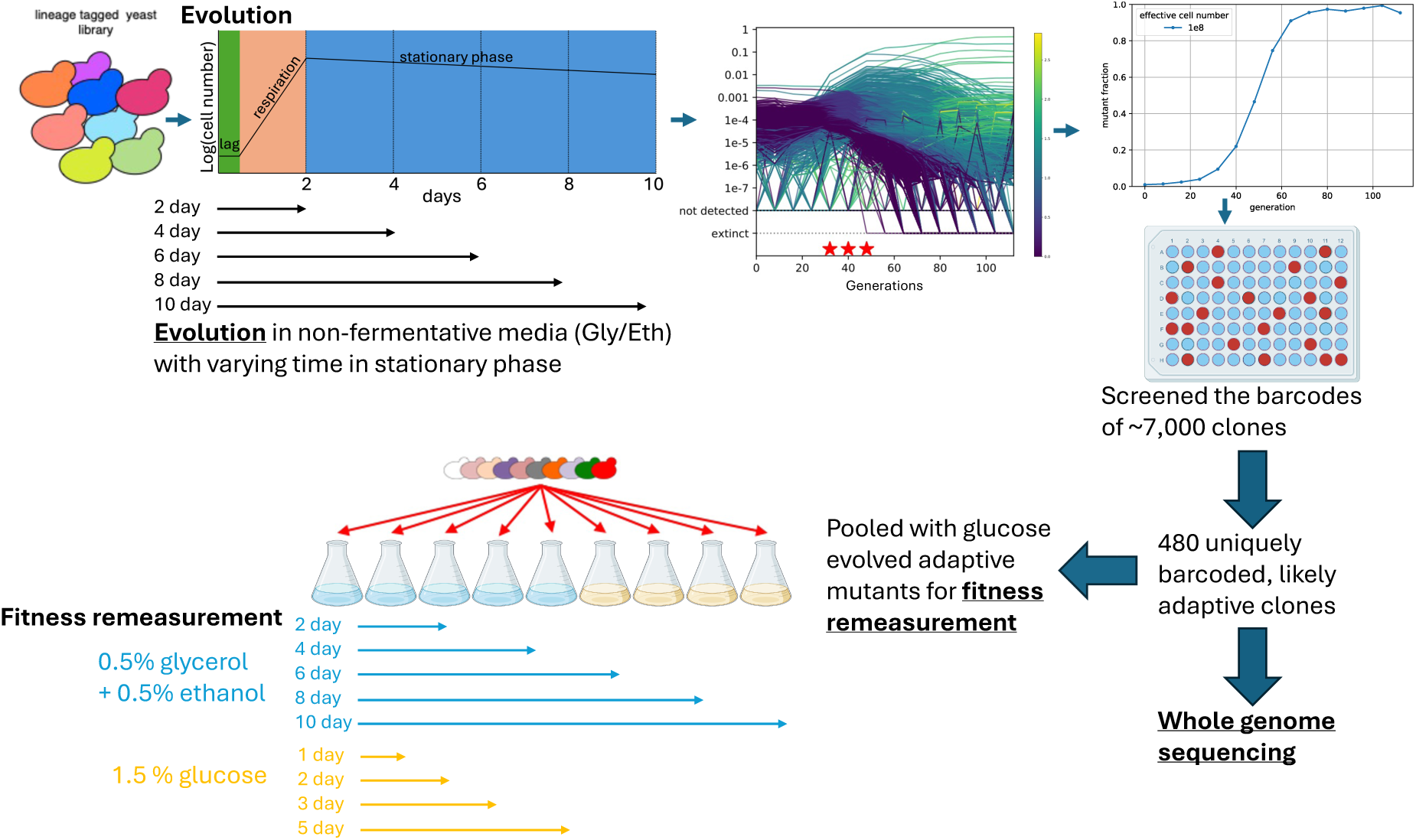
Experimental Design. Barcoded yeast were evolved in triplicate in Gly/Eth medium with 0-8 days spent in stationary phase between transfers. Barcodes were sequenced, counted and used to estimate the adaptive fraction of the population. Clones were screened and 480 uniquely barcoded clones were whole-genome-sequenced, and their mutations were identified. Uniquely barcoded clones were pooled with hundreds of adaptive clones previously evolved in limiting glucose (Levy et al., 2015; Li et al., 2019) and their fitnesses were measured in 9 different conditions.

These clones were then pooled with adaptive clones (Table 1) previously evolved in the presence of the fermentable carbon source glucose (Levy et al., 2015; Y. Li et al., 2019), and their fitnesses *per growth cycle* (defined such that the log fold change over a cycle is the difference between a lineage’s fitness and the population mean fitness) were measured in 9 conditions (Figs. S2, S3, and S4). These fitness measurements allowed us to estimate performance within different parts of the growth cycle, including different intervals of stationary phase following growth in glucose and Gly/Eth (see Table 3 in Methods). Ancestral lineages were included in the fitness assays, allowing us to calculate the *change* in each performance metric for each adaptive clone. We subdivided stationary phase performance into both 2- and 4-day intervals and asked how mutations that change early stationary phase performance affect later stationary phase performance. We also determined whether stationary phase performance is dependent on the type of carbon limitation (Gly/Eth vs. glucose).

**Figure 2.**
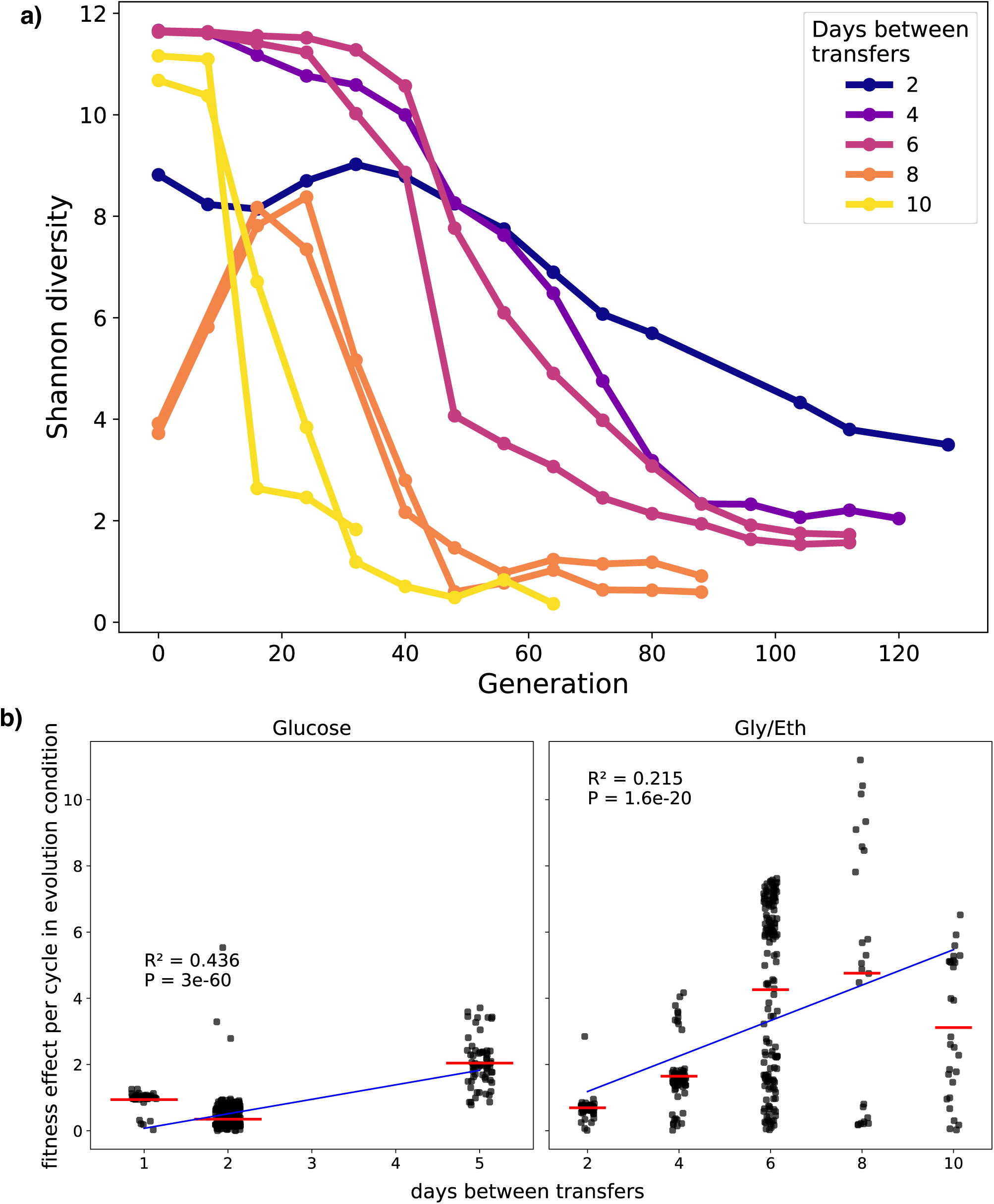
Shannon diversity decreases faster and fitness effects increase with more time spent in stationary phase. (a) Shannon diversity of the barcode pools in each sequenced Gly/Eth evolution replicate over time. (b) Fitness effects of **adaptive mutations** in their evolution condition as a function of transfer time for both glucose (left) and Gly/Eth-evolved clones (right).

**Figure 3.**
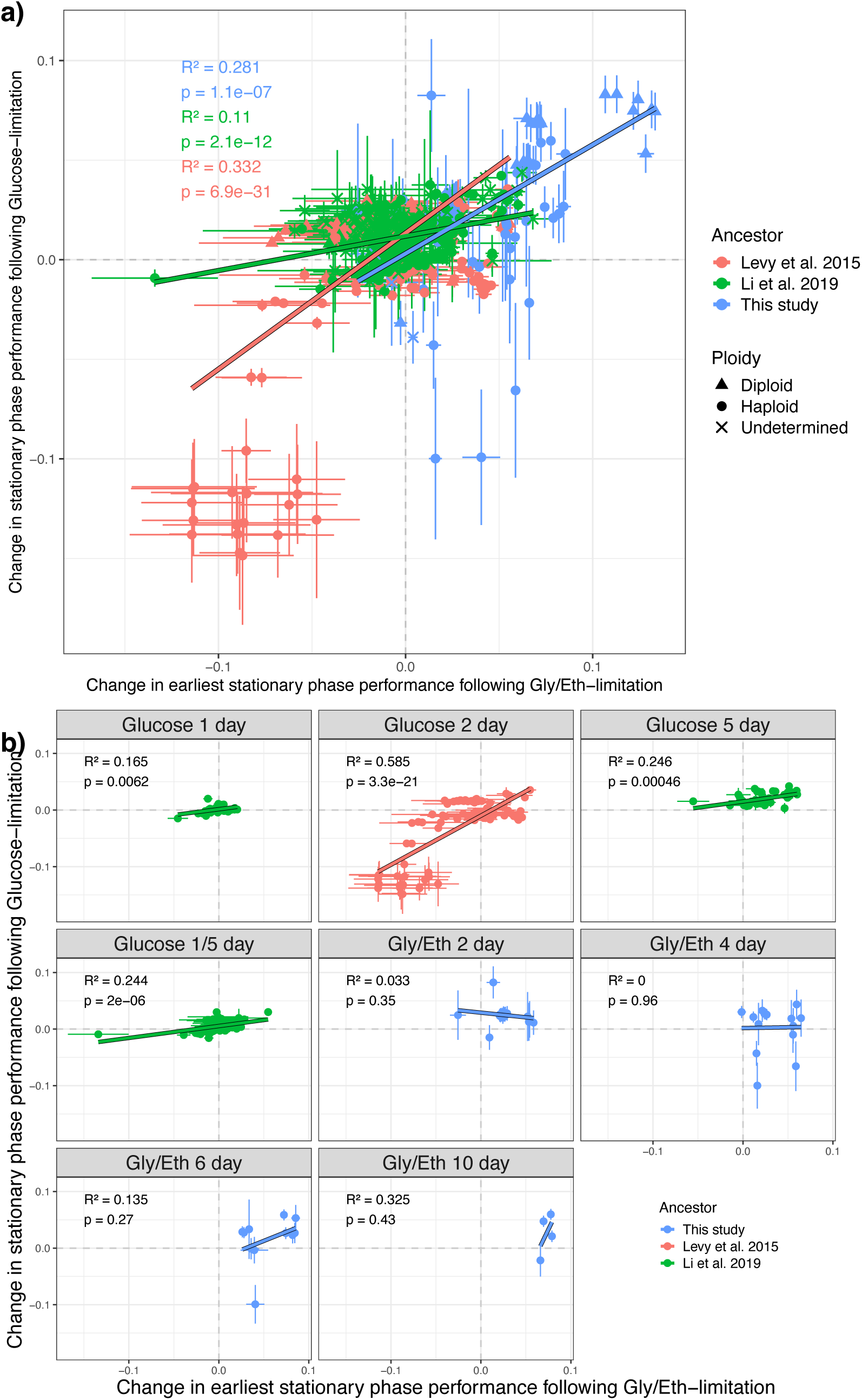
Change in stationary phase performance is correlated following glucose and Gly/Eth-limitation. Changes in stationary phase performance (day 3-5) following glucose-limitation are correlated with changes in earliest stationary phase performance (day 2-4) following Gly/Eth-limitation (Table 3). (a) For each set of <u>all</u> adaptive mutants there is a significant positive correlation between stationary phase performance following glucose- and Gly/Eth-limitation. (b) Different sets of evolved adaptive <u>haploid</u> mutants are shown in each plot. Levy et al. 2015 and Li et al. 2019 clones from each evolution condition all show a strong positive correlation. Error bars indicate the combined error inferred by Fitseq2 for each underlying fitness measurement.

**Table 1.** Evolved clones in our fitness measurement pool derive from 3 ancestors and sets of evolutions. Barcoded clones were previously evolved in glucose with 2 days (Levy et al., 2015) or 1, 5, or alternating 1 and 5 days between transfers (Y. Li et al., 2019) between transfers. Here, we evolved barcoded clones in Gly/Eth medium with 2, 4, 6, 8, or 10 days between transfers.

| Ancestor | Media | Time between transfers |
| --- | --- | --- |
| Levy et al (2015) | Glucose | 2 days |
| Li et al (2019) | Glucose | 1 days |
| Li et al (2019) | Glucose | 5 days |
| Li et al (2019) | Glucose | 1/5 days |
| This study | Gly/Eth | 2 days |
| This study | Gly/Eth | 4 days |
| This study | Gly/Eth | 6 days |
| This study | Gly/Eth | 8 days |
| This study | Gly/Eth | 10 days |

**Table 2.** Beneficial mutations observed from experimental evolution vary depending on time spent in stationary phase. Loci with multiple mutations observed in adaptive clones are summarized above.

| Gene | Pathway/Role | Gly/Eth |  |  |  |  | 5 day gluc. |
| --- | --- | --- | --- | --- | --- | --- | --- |
|  |  | 2 day | 4 day | 6 day | 8 day | 10 day |  |
| *IRA1, IRA2, PDE2, GPB2, YAK1* (see Table. S1 for full list) | Ras/PKA | 60 | 5 | 6 | 1 | 7 | 3 |
| *SSK1* | High osmolarity glycerol (HOG) | 1 | 0 | 0 | 0 | 1 | 7 |
| *SSK2* |  | 3 | 0 | 1 | 0 | 1 | 1 |
| *FPK1* | Ser/Thr protein kinase | 0 | 0 | 0 | 0 | 0 | 10 |
| *SXM1* | Nuclear transport | 0 | 0 | 0 | 0 | 0 | 10 |
| Chr11 duplication | *UTH1*/resistance to ROS | 0 | 4 | 84 | 2 | 14 | 11 |
| YPL150W | Putative Ser/Thr protein kinase; overexpressed in 4% glycerol | 0 | 0 | 3 | 0 | 0 | 1 |
| *IMH1* | Vesicular transport | 0 | 0 | 3 | 0 | 0 | 0 |
| *SMF2* | Metal ion transport | 0 | 0 | 69 | 1 | 7 | 0 |
| *SMF1* |  | 0 | 0 | 1 | 1 | 0 | 0 |
| *CCH1* |  | 0 | 0 | 0 | 2 | 0 | 0 |
| *AIF1* upstream | Mitochondrial cell death effector | 0 | 0 | 0 | 2 | 0 | 0 |
| *FZF1* | TF (transcription factor) involved in sulfite metabolism | 0 | 0 | 0 | 7 | 0 | 0 |
| *ACE2* | TF required for septum destruction after cytokinesis | 0 | 0 | 0 | 0 | 6 | 0 |
| *RAV1* | Regulator of ATPase of vacuoles and endosomes (RAVE) complex component | 0 | 0 | 0 | 0 | 3 | 0 |
| *MPF1* | Unknown function, localized to mitochondria | 0 | 1 | 1 | 0 | 0 | 0 |
| *RPA190* | RNA polymerase I largest subunit | 0 | 1 | 0 | 0 | 1 | 0 |
| *HSL7* | Protein arginine N-methyltransferase | 0 | 1 | 0 | 1 | 0 | 0 |
| No adaptive mutation detected (among clones with sufficient coverage and which were adaptive in fitness remeasurement) |  | 7 | 31 | 7 | 0 | 2 | 0 |
| Adaptive clones with other mutations (SNPs, diploids, and other karyotype changes) |  | 16 | 32 | 8 | 14 | 8 | 29 |
| # of clones in the fitness remeasurement assay (# that were adaptive) |  | 91<br>(78) | 87<br>(76) | 185<br>(152) | 51<br>(24) | 65<br>(28) | 130<br>(124) |
| Clones with sufficient coverage in WGS (see Methods) |  | 86 | 81 | 179 | 49 | 54 | 71 |

**Table 3.** Inferring performance in different parts of the growth cycle from underlying fitness data. Fitness assays in glucose allowed us to infer stationary phase performance between days 3 and 5. Fitness assays performed in Gly/Eth allowed us to infer performance within each 2- or 4-day interval of stationary phase. Herein we consider the earliest (day 2-4), early (day 2-6), late (day 6-10), and latest (day 8-10) stationary phase intervals.

| Performance Metric (hour<sup>-1</sup>) | Definition |
| --- | --- |
| *Stationary phase following Glucose-limitation* | (5 day gluc. fitness – 3 day gluc. fitness)/48 hours |
| *Early stationary phase following Gly/Eth limitation (day 2-6)* | (6 day Gly/Eth fitness – 2 day Gly/Eth fitness)/96 hours |
| *Late stationary phase following Gly/Eth limitation (day 6-10)* | (10 day Gly/Eth fitness – 6 day Gly/Eth fitness)/96 hours |
| *Earliest stationary phase following Gly/Eth limitation (day 2-4)* | (4 day Gly/Eth fitness - 2 day Gly/Eth fitness)/48 hours |
| *Latest stationary phase following Gly/Eth limitation (day 8-10)* | (10 day Gly/Eth fitness – 8 day Gly/Eth fitness)/48 hours |

### Longer periods of stationary phase result in faster loss of barcode diversity, greater fitness effects, and distinct genomic signatures of adaptation

We find that barcode diversity is lost more rapidly in populations whose growth cycles include more time in stationary phase (Fig. 2a). Concomitantly, the fitness effects of adaptive mutations tend to be larger in evolutionary conditions with more time in stationary phase, although this effect appears to saturate for growth cycles with more than 4 days of stationary phase (Fig. 2b). Additionally, distinct classes of mutations are seen to be adaptive in growth cycles with different lengths of stationary phase (Table 2). In clones evolved in Gly/Eth-limited medium with zero days in stationary phase (2 days between transfers), many Ras/PKA mutations are observed (60/87 clones), consistent with prior observations (Chen et al., 2023) and with data from 2-day glucose evolved clones that also experienced minimal stationary phase between transfers (Levy et al., 2015; Venkataram et al., 2016). These Ras/PKA mutants are most often loss of function mutations in negative regulators of the signaling pathway and thus result in increased Ras/PKA signaling. However, with 2 or more days in stationary phase, far fewer Ras/PKA mutants are successful, consistent with previous findings showing a trade-off between the Ras/PKA mutants and stationary phase performance (Y. Li et al., 2019).

While the routes of adaptation were similar for the 2 day evolved clones regardless of the carbon-limiting substrate, clones evolved with stationary phase had distinct routes of adaptation depending on the carbon-limiting substrate. 38% of adaptive clones that evolved with stationary phase following glucose exhaustion evolved via mutations in *SSK1* (7/71), *FPK1* (10/71), and *SXM1* (10/71) (Y. Li et al., 2019). *FPK1* and *SXM1* mutations were not observed in any of the Gly/Eth stationary phase-evolved clones and only 2 *SSK1* mutations were observed among the Gly/Eth clones.

We also saw distinct signatures of adaptation for each additional 2 days of stationary phase per growth cycle. Populations evolved with 2 days in stationary phase (4 days between transfers) tended to have less parallelism than other populations, with the only parallelism observed being Ras/PKA pathway mutations (5/81) and chromosome 11 duplications (4/81). Ras/PKA mutations were primarily observed in clones evolved with 2 days between transfers while chromosome 11 duplications were primarily observed in clones evolved with 6 days between transfers, indicating that the 4-day transfer cycle favors kinds of mutants that are successful in the 2-day or 6-day growth cycles. The lack of observed parallelism amongst the 76 adaptive clones evolved with 4 days between transfers suggests that there are many competing routes of adaptation in this condition.

With 4 days in stationary phase (6 days between transfers) in Gly/Eth there was extensive parallelism: 84.9% of these mutants adapted via (1) a chromosome 11 duplication (46.9%), or (2) an *SMF2* mutation (38.5%). Only one adaptive mutant had both mutations, suggesting possible negative epistasis between the two types of mutations. Indeed, the fitness of the double mutant in the evolutionary condition is lower than all but one of the other *SMF2* mutants, but higher than most of the other chromosome 11 duplications (Figure S5). In total, 77 *SMF2* mutants were found across all 449 clones sequenced with sufficient coverage: 5 of these clones had two mutations in the *SMF2* gene, and 1 had five mutations within the *SMF2* gene. There were 11 positions within the *SMF2* gene that were mutated in multiple clones, accounting for 27/86 *SMF2* mutations. Of these 11 mutated positions, 3 displayed different mutations between clones, indicating that they arose independently and demonstrating parallelism at the nucleotide level.

Clones evolved with 8 days between transfers displayed heterozygous missense mutations in *FZF1* (14.3% of clones). All these clones were diploid, and they were among the fittest clones across all Gly/Eth evolution conditions with a stationary phase (Figure S6). *FZF1* encodes a transcription factor involved in the regulation of sulfite metabolism (Avram et al., 1999; Du et al., 2024; Engle & Fay, 2012; Lin et al., 2023; Park & Bakalinsky, 2000). Clones evolved with 10 days between transfers evolved via mutations in Ras/PKA genes (12.7%), chromosome 11 duplications (25.5%), *SMF2* (12.7%), *ACE2* (10.9%), and *RAV1* (5.45%). The *ACE2* and *RAV1* mutations were observed exclusively in the 10 day evolved populations. *ACE2* is a transcription factor required for asymmetric division of daughter cells and septum destruction after cytokinesis (Herrero et al., 2020; Sbia et al., 2008), and mutations in *ACE2* are known to be involved in the “multicellular” snowflake yeast phenotype previously described in the literature (Ratcliff et al., 2015). Meanwhile *RAV1* is part of the RAVE complex (Seol et al., 2001) which promotes assembly of the V-ATPase holoenzyme and is required for transport between the early endosome and the late endosome/pre-vacuolar compartment (Sipos et al., 2004).

Taken together, our sequencing results indicate that the mutants that are adaptive (with the exception of chromosome 11 duplications, which were observed in evolutions in both fermentable and non-fermentable carbon sources) depend both on the nature of carbon-limitation and the amount of time they spend in stationary phase between transfers.

### Increases in stationary phase performance in glucose and Gly/Eth are correlated

Next, we asked whether the type of carbon-limitation (glucose vs. Gly/Eth) affects the impact of a mutation on performance in the subsequent stationary phase. The cell state adopted in stationary phase may differ as a function of both the carbon source consumed during growth in the spent media, and therefore the same mutant may experience stationary phase differently depending on the carbon source. We found that change in stationary phase performance between days 3 and 5 of the growth cycle in glucose and between days 2 and 4 of the growth cycle in Gly/Eth was significantly positively correlated for each set of evolved mutants that we tested (Figure 3a). Clones evolved under glucose-limitation followed by stationary phase often displayed a pleiotropic increase in Gly/Eth stationary phase performance, while clones evolved with Gly/Eth stationary phase (with more than 6 days between transfers) often displayed a pleiotropic increase in glucose stationary phase performance. These effects were considerable for the glucose-evolved clones while they were not significant for the Gly/Eth clones from this study (Figure 3b), perhaps as a result of the smaller number of Gly/Eth clones from each evolution condition that were accurately assayed (see Methods) in the glucose conditions. If we include diploids in our analysis, we observe a positive correlation between change in stationary phase performance following Gly/Eth and glucose limitation amongst clones from all conditions that evolved with stationary phase, and a significant positive correlation for the Gly/Eth 8 and 10 day evolved clones (Figure S7). Overall mutational effects on stationary phase performance were similar regardless of the carbon source limitation.

### A trade-off exists between early and late stationary phase performance

We compared the mutational effect on performance during the first 2 days of stationary phase (earliest SP, day 2-4) to the mutational effect on performance during the last 4 days of stationary phase (late SP, day 6-10; Figure 4). We observed significant negative correlations among all adaptive clones evolved from each ancestor (Figure 4a). We also asked whether the specific evolution condition affected the relationship between change in early and late stationary phase performance. Among adaptive haploid clones we found a negative correlation between change in early and late stationary phase performance for 4- and 10- day Gly/Eth mutants but not for the 2- and 6-day Gly/Eth mutants (Figure 4b). The 2-day Gly/Eth clones did not experience stationary phase selection and thus show minimal change in early stationary phase performance. The 6-day Gly/Eth clones have a positive relationship (Figure 4b) because of the *SMF2* mutations that arise in this condition. However, for the 6-day evolved Gly/Eth clones, though the *SMF2* mutants’ change in performances cluster, indicating reliability of our measurements, a negative correlation is observed between earliest and late stationary phase for clones within this cluster (Figure S8).

**Figure 4.**
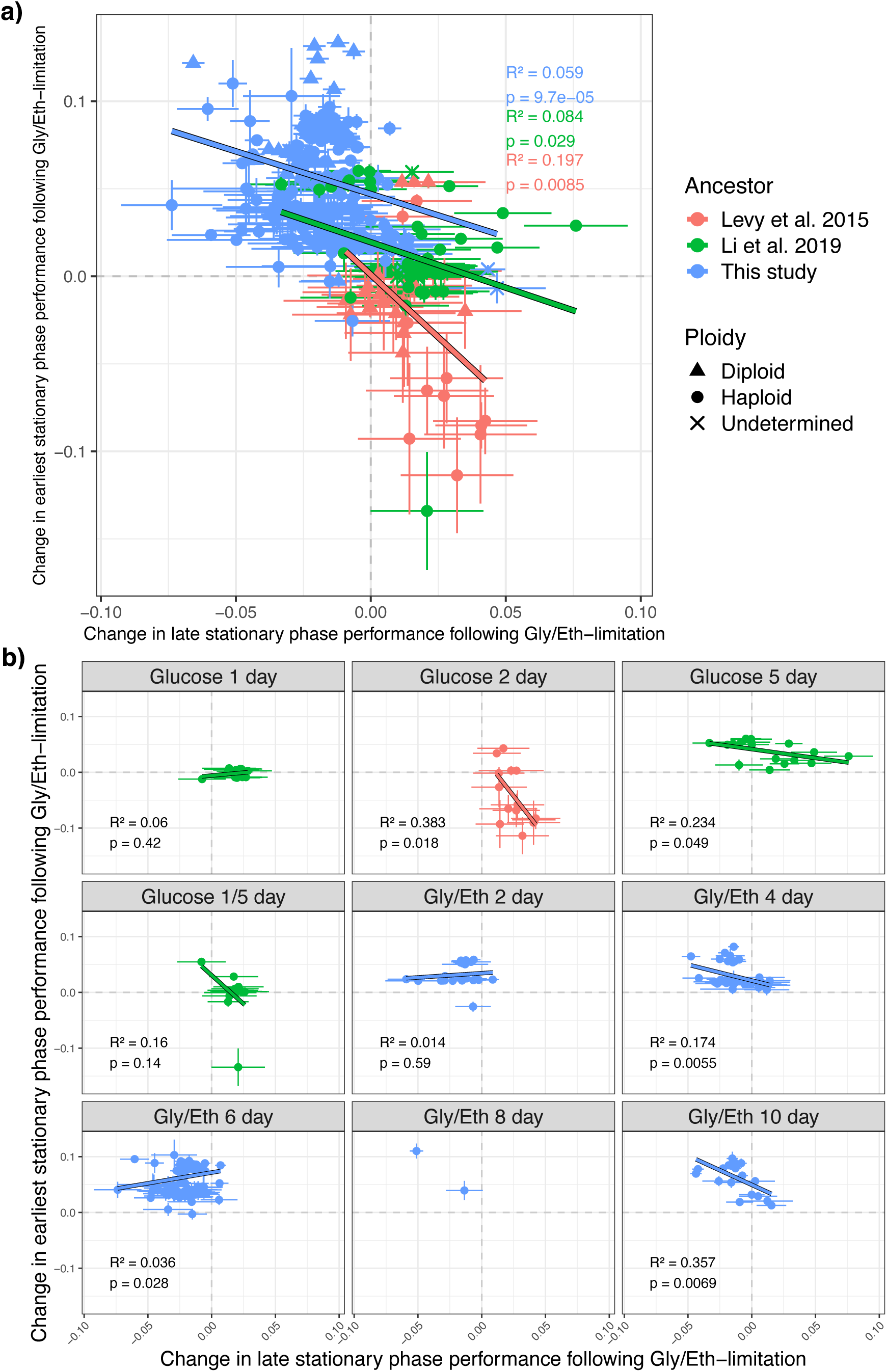
Performances in early and late stationary phase show evidence of a trade-off. Changes in late stationary phase performance (day 6-10) following Gly/Eth-limitation are negatively correlated with changes in <u>earliest</u> stationary phase performance (day 2-4) following Gly/Eth-limitation. (a) For each set of <u>all</u> evolved mutants there is a significant positive correlation between change in late and earliest stationary phase performance following Gly/Eth-limitation. (b) Different sets of evolved adaptive <u>haploid</u> mutants are shown in each plot. All evolved conditions that include a stationary phase show a negative correlation between early and late stationary phase performance except for the Gly/Eth 6 day clones. However, the Gly/Eth 6 day clones are negatively correlated if *SMF2* mutants, which are 38.5% of these clones and have large fitness effects, are treated separately (Figure S6). Error bars indicate the combined error inferred by Fitseq2 for each underlying fitness measurement.

We also compared performance during the first 4 days of stationary phase (early SP, day 2-6) to the last 2 days of stationary phase (latest SP, day 8-10; Figure S9a-b) and saw similar negative correlations between early and latest stationary phase performance. We purposefully avoided using the same underlying fitness data in the calculation of the performance metrics we compared (e.g., we do not directly compare performance in early SP and late SP because the Gly/Eth 6 day fitness is used in both performance metrics). When we removed diploids and *SMF2* mutants (known to have significant phenotypic effects) from both of these analyses (Figure 4a and S9a), the proportion of the variance in early stationary phase that is explained by performance in late stationary phase increases (Figure S10a-b), indicating the trend is not driven by the effect of *SMF2* mutations or diploidization. These findings demonstrate a trade-off between early and late stationary phase performance and indicate that stationary phase performance is affected more by time in stationary phase than by the type of carbon limitation. A clone that has high performance in stationary phase following glucose-limitation is likely to perform well following Gly/Eth-limitation, while a clone with high early stationary phase performance is likely to have low late stationary phase performance.

### Changes in early and late stationary phase performance trade-off within a single gene

We isolated 77 mutants from the Gly/Eth evolutions that contained *SMF2* mutations. These included a mix of frameshift, nonsense, and missense mutations found throughout the gene. The *SMF2* mutants cluster together in trait-space separately from other evolved mutants (Figure S8). All *SMF2* mutants have substantial increases in early stationary phase performance compared to their ancestor, and all but one show a decrease in late stationary phase performance, indicating a potential trade-off in late stationary phase performance. Haploid *SMF2* mutants with normal karyotypes (n = 76) showed a notable negative correlation between their mutational effects in early and late stationary phase, though this correlation is not easily explainable by the type of mutation that caused the adaptation (Figure 5 and S11). For the *SMF2* mutants, a large proportion of the variance in early stationary phase performance can be predicted by late stationary phase performance (33-47%, Figure 5 and S11).

**Figure 5.**
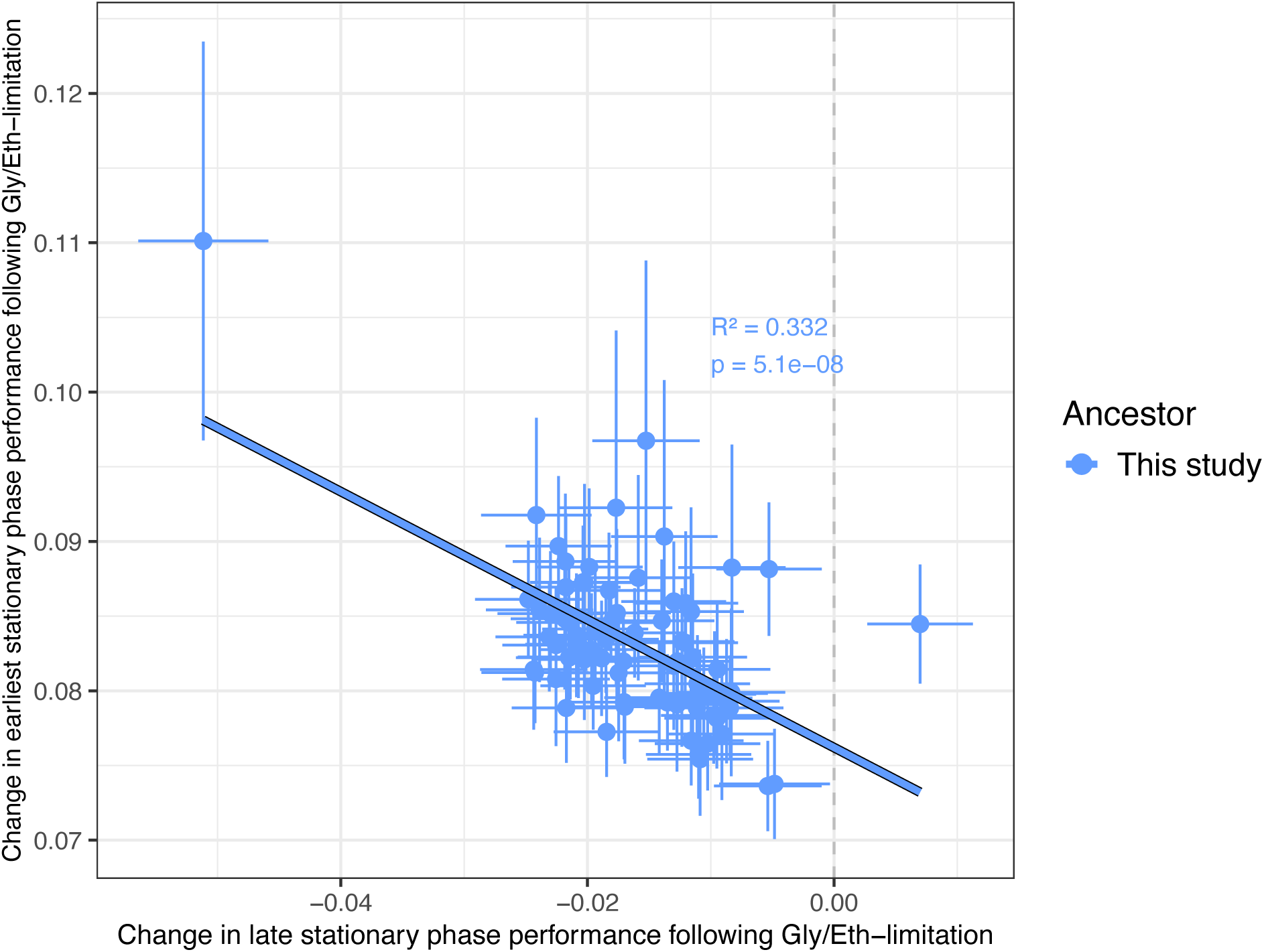
Trade-offs between early and late stationary phase performance among karyotypically normal, haploid *SMF2* mutants. Changes in <u>earliest</u> stationary phase performance (day 2-4) following Gly/Eth-limitation are significantly negatively correlated with changes in <u>late</u> stationary phase performance (day 6-10) following Gly/Eth limitation. Error bars indicate the combined error inferred by Fitseq2 for each underlying fitness measurement.

## Discussion

By studying the effect of stationary phase on evolutionary dynamics in barcoded *S. cerevisiae* populations, we have made a number of findings:

### Longer periods of stationary phase result in faster loss of barcode diversity, greater fitness effects

We found that populations of barcoded yeast evolving with longer periods of stationary phase between transfers lost barcode diversity more quickly and incurred mutations with larger effect sizes, perhaps due to the prolonged stationary phase allowing the mutant advantage over the wildtype to accrue further. Additionally, fitness-mediated epistasis suggests that the effects of mutations tend to be larger in less fit backgrounds (Bakerlee et al., 2022; Reddy & Desai, 2021). If the ancestral population is poorly adapted to an environment with more time in stationary phase, such epistasis would predict a larger adaptive mutational effect than if the ancestor were well-adapted. Such larger mutations spread more quickly through the population, purging barcode diversity in the process. Our observation that the stationary phase-adapted mutants have large fitness effects is consistent with the large effect size previously observed in bacterial stationary phase cultures where GASP mutants (in *E. coli*) have large increases in frequency within a single growth cycle (Zambrano et al., 1993).

### Distinct genomic signatures of adaptation depending on carbon substrate and time in stationary phase

We observed distinct patterns of parallelism in the genomic adaptations which arose during evolution, depending on the carbon source and on the amount of time between serial dilutions. The most common parallel mutation, which was found at high frequency in the Gly/Eth 6 day populations, was in the *SMF2* gene. *SMF2* encodes an intracellular ion transporter involved in manganese homeostasis (Cohen et al., 2000; Portnoy et al., 2002). Elevated manganese levels are also known to increase TOR signaling (Nicastro et al., 2022), so it is possible that these mutations affect TOR signaling pathways. We also observed 3 *YPL150W* mutations in the 6-day evolved condition. As *YPL150W* deletion strains form fewer TORC1 bodies during starvation (Sullivan et al., 2019), these mutations may act to maintain TOR signaling during starvation, perhaps in anticipation of the next transfer to avoid entering a quiescent state and thus reducing the lag time in the next growth cycle.

We also observed a number of chromosome 11 duplications, which have previously been shown in yeast to be adaptive in the presence of the toxin paraquat (Helsen, 2021), which produces reactive oxygen species (ROS) (Fukushima et al., 2002). Chromosome 11 contains the *UTH1* gene which is known to be involved in the oxidative-stress response (Bandara et al., 1998), and increasing expression of *UTH1* results in a similar paraquat-resistant phenotype as the chromosome 11 duplications (Helsen, 2021), suggesting that increasing *UTH1* copy number may be why the chromosome 11 duplication is beneficial in our fitness measurement assays. Duplicating the entire chromosome may be the easiest way to increase *UTH1* copy number and any deleterious effect of this karyotype change might be outweighed by the benefit of increased expression of *UTH1*. *UTH1* mutants have also previously been observed to arise in response to ethanol selection (Avrahami-Moyal et al., 2012) and additional work has implicated chromosome 11 duplications as an adaptation in response to oxidative stress (Kaya et al., 2015). In our fitness measurement assays, the chromosome 11 duplications are consistently beneficial in the 6-day condition in which they arose and have smaller fitness gains in the 2-day, 4-day, and 10-day Gly/Eth condition (Figure S12), which may be due to the transient nature of ROS in stationary phase cultures.

ROS build up during the proliferative portion of the growth cycle as a byproduct of oxygen metabolism (Krumova & Cosa, 2016; Pryor, 1986). After this point, new ROS are no longer being produced via respiration, and the ROS present will begin to diffuse and break down into less reactive species (Winterbourn, 2008). As a result, ROS stress could be maximal in the 6-day transfer regime. In the 4-day condition, selection for growth-related traits may dominate, while in the 8- and 10-day conditions ROS may be less prevalent at the end of stationary phase and other types of adaptations might be favored. These results indicate that the genomic signature of adaptation depends on both the carbon-limitation and the amount of time spent in stationary phase.

We note that the library used to found the 8-day Gly/Eth populations had less barcode diversity at the beginning of the experiment. This low diversity is likely an artifact of the transformation process which made it difficult to isolate unique adaptive clones from these populations and contributed to the smaller number of adaptive mutants isolated from this condition.

### Increases in stationary phase performance in glucose and Gly/Eth are correlated

We also assessed whether the carbon source in which yeast grow impacts the effect of mutations during the subsequent stationary phase (Galdieri et al., 2010). While the mutations which arose in glucose vs Gly/Eth evolutions with similar time spent in stationary phase (e.g., 4 days) were different, the performance of all adaptive mutants was roughly similar in stationary phase following glucose exhaustion and in stationary phase following Gly/Eth exhaustion. This suggests that the differences in the types of mutations observed may result from differences in the proliferative portion of the growth cycle (fermentation vs. respiration). Across the different types of mutations and carbon sources investigated, mutational effects in stationary phase were similar.

### A trade-off exists between early and late stationary phase performance

In contrast to the positive correlation observed between stationary phase performance following glucose vs. Gly/Eth exhaustion, we observed a negative correlation between early and late stationary phase performance. In fact, we observed that the mutations that are most beneficial in early stationary phase often have a deleterious effect in late stationary phase, and *vice versa* (Figure 4). Interestingly, we also observed this trade-off within a single gene. *SMF2* was the most frequent mutational target observed in this study, which can be explained in part because these mutants tend to have high early stationary phase performance.

### Outlook and Further Questions

The population dynamics in stationary phase yeast cultures differ from those observed in bacterial cultures (Shoemaker et al., 2021; Zambrano et al., 1993). Bacteria persist for long periods in stationary phase culture, aided by successive mutations that often sweep through the population (Zambrano & Kolter, 1996). Yeast, on the other hand, has a finite chronological lifespan with some variability (Fabrizio & Longo, 2003). Part of this variability in lifespan may result from the occurrence of GASP-like mutants that grow in the spent media and extend the life of the culture (Garay et al., 2014). When rare GASP mutants are observed in the lab, they do not appear to occur in succession and only extend the lifespan temporally (Fabrizio et al., 2004). Compared to bacteria, yeast cells may be more sensitive to the stresses of stationary phase culture, making their lifespan short and mutations that allow growth in stationary phase rare. In addition, the smaller population sizes in yeast experiments compared to bacteria (10-100 times smaller (Fabrizio et al., 2004; Zambrano et al., 1993)) make these mutations even less likely to occur within a single growth cycle, whereas they occur systematically in stationary phase cultures of *E. coli.* However, the evolutionary dynamics of yeast in even longer periods of stationary phase or in larger populations remain to be explored.

One of our main findings is that stationary phase in yeast is dynamic, with mutations having negatively correlated effects on change in performance in early and late stationary phases. This was consistent for clones from the three genetic backgrounds included in our fitness measurements. This conclusion is based on an analysis which isolates the component of fitness derived from a part of the growth cycle (e.g., days 8-10 of the cycle) by subtracting the fitness without this part of the cycle from the fitness with this part of the cycle included. Although this definition of performance is a reasonable heuristic, it neglects the fact that extending the growth cycle does more than simply allow for the accrual of a different set of fitness benefits: it also changes the state of the cells when they are diluted into fresh media, making days 1-8 of the 10-day cycle different from days 1-8 of the 8-day cycle. Measurements during a growth cycle, as performed in Li et al 2018, would further support our claims about the dynamism of stationary phase, and it is possible that such finer resolution measurements would elucidate further tradeoffs between mutational effects during different parts of stationary phase. Additionally, extending stationary phase further may reveal whether performance after 10 days is the same or distinct from late stationary phase performance assessed in this study (day 6-10).

In nutrient-limited environments, constraints on performance in different intervals of stationary phase could play an important role in shaping microbial diversity and guiding life-history evolution. Future investigations should explore whether the observed trade-off between early and late stationary phase performance is a common feature of microbial systems or specific to the *Saccharomyces cerevisiae* strains used in this experiment.

## Methods

### Barcoded Yeast Libraries

We used previously-constructed haploid barcoded yeast pools (Boyer et al., 2021; Chen et al., 2023): pools 1 (10-day populations), 3 (8-day populations), 5 (6-day populations), 6 (4-day populations), and 7 (2-day populations) were used in our experiments.

### Growth Media

Cells were grown in 5x Delft with 4% ammonium sulfate supplemented with vitamins and minerals, as previously described (Verduyn et al., 1992), with either 1.5% glucose (glucose medium; Levy et al. 2015; Venkataram et al. 2016; Li et al. 2019; Kinsler et al. 2024) or 0.5% ethanol and 0.5% glycerol (Gly/Eth medium; this study).

### Growth Curve Measurement

The growth curve of our barcoded library in Gly/Eth medium was measured in 2 staggered growth curve experiments (Figure S13). Cells from pool 1 were pre-cultured overnight in 100 mL of Gly/Eth medium at 30°C with shaking at 200 RPM. The following day, ∼3.6e7 cells were inoculated into in 100 mL of Gly/Eth medium and incubated at 30°C while shaking at 200 RPM. Measurements were taken immediately following inoculation and then again early the next morning (16 hours later) and approximately every 2 hours during daytime hours. Measurements included OD600, a Coulter Counter reading, and CFUs/mL. In the morning after the first inoculation another replicate culture was independently established by inoculating ∼5.6e7 cells into a new 100 mL of Gly/Eth medium that was incubated at 30°C while shaking at 200 RPM. Measurements were also taken on this culture approximately every 2 hours.

### Experimental Evolution

All evolution experiments were started by inoculating at least 1.4e8 cells (there was some variability in the number of cells in each frozen aliquot) from one of the barcode pools into 100 mL of 5x glucose medium (Levy et al., 2015; Verduyn et al., 1992). Cells were grown in non-baffled flasks while incubating at 30°C and shaking at 200 RPM. The following day, 0.6 mL of culture was used to inoculate 2-3 replicate cultures (2 replicates for the 2-day populations and 3 replicates for the others) each containing 100 mL of Gly/Eth media (Verduyn et al., 1992). This transfer volume was adjusted slightly (±0.1 mL) depending on the Coulter Counter reading to keep the total number of cells transferred standardized across experiment. Cultures were allowed to incubate at 30°C while shaking at 200 RPM for 2, 4, 6, 8, or 10 days. Subsequent transfer volumes were adjusted for the number of viable cells (measured as CFUs/mL from plating) from the previous timepoint so that ∼5e7 cells (0.4 – 1.0 mL) were transferred at each timepoint. Populations were allowed to evolve in this manner for between 16-25 transfers or the equivalent of 128-200 generations.

At each transfer, an OD reading and Coulter Counter reading were obtained and ∼200 cells were plated onto 2 YPD plates and 2 URA- plates to monitor for viability and contamination. The barcoded yeast strain used in the evolutions can grow on URA-plates while many common laboratory contaminants cannot. After each transfer, 5 mL of leftover culture were removed, centrifuged, and resuspended in 2 mL of 17% glycerol.

Five 400 µL aliquots were stored at -80°C. The remaining 95 mL of culture were divided into four 50 mL conical tubes, centrifuged, washed once with 2 mL of TE buffer, and then resuspended in 1 mL of 0.9M sorbitol (0.9M Sorbitol, 0.1M Tris-HCL, pH7.5 0.1M EDTA, pH8.0) and stored at -20°C. At each timepoint ∼96 clones were isolated from the URA- plates (or YPD plates if there were not enough colonies on the URA- plates) into 96-well plates. Plates were allowed to grow to saturation before being frogged onto benomyl containing medium for ploidy determination (Venkataram et al., 2016), after which glycerol was added, and plates were duplicated and then stored at -80°C.

### Ploidy Determination

Ploidy was determined in real time via benomyl assays (Figure S14), as described in Venkataram et al., 2016. Ploidy was verified via FACS for the set of clones that were whole genome sequenced using a modified version of the protocol described in Todd et al. 2018 (instead of sonication, plates were vortexed to resuspend cells).

### DNA extraction for Barcode Amplification

We developed a high-yield method for extracting high-purity DNA from yeast cells. The frozen pellets (from ∼24 mL of culture) in sorbitol were allowed to thaw to room temperature. Samples were centrifuged at 2,900 RPM and the supernatant was discarded. Pellets were resuspended in X buffer (1.4M NaCl, 2% PVP, 2% CTAB, 100 µM Tris, 10 µm EDTA) and transferred to an O-ring tube containing 250-300 µL of beads. 4 µL of RNAse were added and the cells were bead beaten for 5 minutes. The tube was then incubated at 65°C for 30 minutes. After this, 400 µL of chloroform:isoamyl (24:1) were added and the tube was vortexed (30 seconds) followed by another 15-minute incubation at 65°C. The tube was then centrifuged on high for 5 minutes and the top layer (containing the DNA) was moved to a new Eppendorf tube. 600 µL of chloroform:isoamyl (24:1) were added and the tube was vortexed (5 seconds) then centrifuged at 15,000 RPM for 2 minutes. The top layer was again moved to a new Eppendorf tube; this step was repeated a third time if necessary (the interphase was still thick). The tube was then split into two tubes if necessary and isopropanol was added (70% of volume of sample) to each tube. The sample was then mixed gently and centrifuged at 15,000 RPM for 5 minutes before removing the supernatant. 1 mL of 70% ethanol was then added to the sample. Tubes were then centrifuged at 15,000 RPM for 30s and all supernatant was removed, and the pellet was allowed to air dry. Next, the pellet(s) were resuspended in 200 µL of dH_2_0 (across 1 or 2 tubes) and vortexed thoroughly and samples that were split were repooled. 200 µL of QX1 buffer (Qiagen) were then added and the yellow color was verified (if not yellow, pH was adjusted with 3M sodium acetate). The tube was then vortexed for 30s and again centrifuged at high speed. The supernatant (containing the DNA) was then moved to a new tube. 20 µL of Qiaex2 (from Qiagen, ID 20021; per 10 µg of DNA expected) were then added to each sample and tubes were incubated at room temperature for 10 minutes while mixing every 2 minutes to keep silica beads suspended. The tubes were then centrifuged at 15,000 RPM for 30s and the supernatant was removed. The pellet was then washed twice with 500 µL of PE buffer and resuspended in PE buffer (Qiagen) during the last wash. The pellet was then air dried for 10-15 minutes or until it turned white with care taken not to over dry. 20 µL of 10 µM Tris-Cl were then added back to the tubes and the pellet was resuspended by vortexing. Next, tubes were incubated at 50°C for 10 minutes before being centrifuged at 15,000 RPM for 30 seconds. Finally, the supernatant was transferred to a new Eppendorf tube, and the DNA concentration was measured on the Qubit.

### Barcode Amplification

Barcodes were amplified using a modified version of the 2-step directed PCR described in Levy et al. 2015. The procedure was modified as described below to use indexed primers in the 2^nd^ PCR reaction. Use of dual unique indexing, in which amplicons from each sample have uniquely indexed ends, allowed us to detect index hopping in these samples.

#### Part 1

The first step of a 2-part PCR was performed. 4 µg of input DNA was used across 6 PCR reactions (0.75 µg/tube). Reactions were set up on ice and the PCR block was prewarmed.

PCR reaction components:

25 µL – 2x OneTaq

0.5 µL – 10 µM forward primer (F2**)

0.5 µL – 10 µM reverse primer (R3**)

2 µL – 25 µM MgCl

(X) µL – DNA (0.75 µg)

(22 µL – X) µL - dH_2_0

PCR protocol:

1. 94°C 10 min

2. 94°C 3 min

3. 55°C 1 min

4. 68°C 1 min

5. Repeat steps 2-4 4 times

6. 68°C 1 min

7. Hold at 4°C

#### Part 2

PCR reactions from the same sample were pooled into a single Eppendorf tube. 0.1x the volume of the pooled sample of 3M sodium acetate and 0.7x the volume of isopropanol was added to each tube. Tubes were then centrifuged at 15,000 RPM for 10 minutes. The supernatant was removed, and the pellet and tube were washed with 500 µL of 80% ethanol. The samples were then centrifuged at 15,000 RPM for 30 seconds and all supernatant removed. The tube was centrifuged again at 15,000 RPM and residual supernatant was removed. The tube was then allowed to air dry for 10 minutes.

#### Part 3

The pellet was resuspended in 25 µL of dH_2_O. 50 µL of room temperature Ampure XP beads (from Beckman) were added and mixed by pipette 10 times. Samples were then incubated at room temperature for 10 minutes then mixed again. Tubes were then placed on magnetic racks and beads were allowed to fractionate (∼2 minutes). The supernatant was removed, and the bead pellet and tube were washed with 0.5 mL 80% ethanol for at least 30 seconds. The wash step was repeated once and supernatant removed before the tube was centrifuged, put back on magnet, and all liquid was removed from the tube with a pipette. The beads were then immediately resuspended in 20 µL water and incubated at room temperature for 5 minutes. 11 µL (0.55x volumes) of 20% PEG 8000 2.5M NaCl solution were then added to each sample, and tubes were mixed well and incubated for 10 minutes at room temperature. Tubes were mixed again, and the beads were allowed to separate on the magnet. The supernatant (containing the DNA) was aspirated to a new tube and 77.5 µL (2.5x volumes) of 100% ethanol were added. Samples were then put on ice for 10 minutes, then spun at 15,000 RPM for 10 minutes. The supernatant was removed, and the DNA pellet was washed with 500 µL of 80% ethanol. The tubes were spun again, and any remaining liquid was completely removed. Tubes were then left to air dry for 10 minutes. The DNA pellet was then resuspended in 20 µL of EB and kept on ice. DNA concentrations were verified by measuring 1 µL of each sample with the Qubit DNA kit.

#### Part 4

The second step of the 2-part PCR was performed. Reactions were set up on ice and the PCR block was prewarmed.

PCR reaction components:

25 µL of Q5 DNA polymerase

10 µL of 1st part library DNA

2 µL of S5**

2 µL of N7**

11 µL dH_2_0

Baseline (pre-PCR) DNA concentrations were checked on the Qubit. PCR protocol:

1. 98°C 30s

2. 98°C 10s

3. 62°C 20s

4. 72°C 30s

5. Repeat steps 2-4 15 times

6. 72°C 3min

7. Hold at 10°C

DNA concentration was checked post-PCR on the Qubit to ensure amplification. Additional PCR cycles were added as necessary on insufficiently amplified samples.

#### Part 5

Each 50 µL PCR reaction was run on a 2% agarose gel. The barcode band at ∼300bp was excised from the gel and extracted using the QIAquick Gel Extraction Kit (Qiagen, ID 28704).

### Construction of Lineage Trajectories and Analysis Using FitMut2

Amplicon libraries from the evolutions were mixed with genomic DNA samples and sequenced on Illumina’s HiSeq platform. The data were demultiplexed based on the 1^st^ step PCR primers (F and R primers; data were received demultiplexed based on the 2^nd^ step N and S primers). This resulted in barcode reads that are grouped by the primer indices which indicate the experiment and timepoint from which the barcodes were extracted. Barcodes were then dereplicated based on their UMI (unique molecular identifier). A sufficient number of time points from at least 1 replicate population was sequenced to identify timepoints from which to select adaptive clones.

Bartender was used (Zhao et al., 2018) to cluster and count barcodes in each experimental timepoint. FitMut2 (Li et al., 2023) was then used to infer fitness and the probability that a given lineage contains a beneficial mutation at a given timepoint.

### Checking the Barcodes of Isolated Clones

#### Isolating clones

Timepoints were selected where FitMut2 confidently estimated that 30-40% of the population contained beneficial mutations (Figure S1). Earlier timepoints were selected for the 8-day populations due to insufficient barcode diversity in the later timepoints.

Plates containing clones collected during the experiments on which benomyl assays were performed were removed from -80°C storage, and cells were frogged into a new 96-well plate containing YPD. Glycerol stocks of evolved populations from the same timepoints were also removed from the freezer and cells were plated on YPD to isolate additional clones. These plates were incubated at 30°C for 2-3 days, after which colonies were picked into individual wells of a 96-well plate. Cells were grown to saturation and clones were stored in glycerol at -80°C.

#### Lysing isolated clones for barcode identification and whole genome sequencing

PCR lysis buffer (Kwiatkowski Jr et al., 1990) was prepared (excluding MgCl_2_ and gelatin). Yeast clones were inoculated into 100 µL of YPD in individual wells of a 96-well plate and allowed to grow to saturation overnight at 30°C. The next day, the plate was spun down for 3 minutes at 3,000 RPM and the supernatant was removed. 30 µL of lysis buffer were added to each well and the cells were resuspended and transferred to a new 96-well PCR plate. The plate was incubated for 1 hour at 37°C followed by 10 minutes at 95°C. The plate was then submerged in liquid nitrogen or ethanol with dry ice for 2 minutes to flash freeze. Plates were stored at -80°C prior to PCR.

#### Grid-Array PCR

All clones underwent PCR in a grid array (modified from Hays et al. 2023) and barcode amplicons were sequenced in two batches. We used 72 unique forward primers and 64 unique reverse primers which target the DNA barcode and allow us to uniquely index 4608 (72 x 64) samples for sequencing.

Forward primers were prepared by diluting 4 µL of 10 µM forward primers into 234 µL of OneTaq 2x master mix. Reverse primers were prepared by diluting 18 µL of 10 µM reverse primer into 702 µL of dH_2_O. 12.5 µL of diluted reverse primer and 10 µL of diluted forward primer were added to each well of a 96-well PCR plate. 5 µL of lysed cells were then added to each well of the plate. The reaction was prepared on ice and the PCR block was pre-warmed.

PCR protocol:

3 minutes at 94°C

40 cycles of

20 sec at 94°C

30 sec at 48°C

30 sec at 68°C

Hold 4°C

After the PCR, 5 µL from each well were combined and the concentration of DNA was measured using a Qubit for each plate. In total, 72 plates were processed in this way for a total of 6,912 clones. Up to 6 plates worth of clones were then combined based on the Qubit reading to ensure an approximately equal amount of DNA across plates. 20 µL of pooled DNA was then run on an e-gel (Invitrogen, #G700802) to purify. The DNA from the e-gel was diluted 1:500 in dH_2_O and a second PCR reaction was prepared.

PCR reaction components:

10 µL of Q5

2.5 µL of PE1

2.5 µL of PE2

3 µL of dH_2_0

2 µL of 1:500 dil DNA

PCR protocol:

2 min 98°C

12 cycles

10s 98°C

20s 60°C

15s 72°C

Hold 4°C

After the PCR reaction, the cleanup was performed using the QIAquick PCR Purification Kit (Qiagen, #28104), and the DNA concentration was checked on the Qubit. Samples were then mixed so that an equal amount of DNA from each sample was included, and the pooled sample was sent for sequencing.

### Analyzing Barcode Sequences and Selecting Clones for WGS

Reads were first demultiplexed by their unique combinations of F and R primer indices and the consensus barcode was determined. These barcodes were then cross-referenced to the lineage trajectory data, and an inferred fitness and probability of containing a beneficial mutation were assigned to each clone (Figure S1). The clones with the greatest chance of containing beneficial mutations (480) were then selected for whole genome sequencing and re-arrayed into new 96-well plates. The same 480 clones were also pooled for fitness measurement.

### Whole Genome Sequencing

Clones were lysed in 96-well plates as described above. The tagmentation reactions were performed as follows in a modified version of the Illumina-recommended protocol (Kryazhimskiy et al., 2014). TDE1 was mixed with TD buffer at a 1:5 ratio and 1.5 µL were added to each well of a 96-well plate. 1 µL of lysed cells was transferred to each well of the 96-well plate and the plate was incubated at 55°C for 5 minutes.

We used a set of 24 reverse (N) primers and 16 forward (S) primers to amplify and uniquely index DNA from each clone. In this way 384 clones could be sequenced in a single lane and the unique combination of N and S primers could be used to identify each genome. PCR reactions were prepared in 96-well plates. Row master mixes were prepared by adding 23 µL of 2x Kapa to each well of an 8-well strip and then adding 7.65 µL of the required S primer to each well. 2.5 µL were added to each well of the plate so that each row contained the same S primer. Column master mixes were prepared by adding 15.34 µL of 2x Kapa to each well of a 12-well strip and then adding 5.1 µL of the required N primer to each well. 2.5 µL of each diluted N primer were then added to the plate so that each column contains the same N primer. The PCR reactions were prepared on ice and the PCR block was pre-warmed. The PCR reaction was run as follows:

72°C for 3 min

98°C for 2:45 min

98°C for 15 sec

62°C for 30 sec

72°C for 1:30 min

Repeat steps (3–5) 12 times

Hold at 4°C

Reconditioning PCR master mix was prepared by diluting 49 µL of primer P1 and 49 µL of primer P2 into 832 µL of 2x Kapa. 9.5 µL of the master mix were then added to each well of the 96-well plate. This was done on ice and the PCR block was pre-warmed. The reconditioning PCR was run as follows:

95°C for 5 min

98°C for 20 sec

62°C for 20 sec

72°C for 30 sec

Repeat steps (2-4) 4 times

72°C for 2 min

Hold at 4°C

17 µL of room temperature Ampure XP (Beckman Coulter, #A63881) beads were added to each reaction. The plate was then incubated at room temperature for 5 minutes. The plate was then put on the magnetic stand and the beads were washed twice with ethanol following the protocol from Beckman Coulter. After removing the last wash, the plate was allowed to air dry for approximately 5 minutes, taking care not to over dry the beads. The DNA was eluted using 30 µL of EB buffer. The DNA concentration of 8 diagonal wells across each plate were checked on the Qubit to ensure DNA was amplified evenly across clones. 5 µL from each well were then combined for sequencing on the NovaSeq platform.

### Analysis of Whole Genomes

480 adaptive clones and 96 ancestors were sequenced. The reads were aligned to the S288C reference genome R64-3-1_20210421 using bwa (H. Li, 2013), bam files were created, and mutations were called using the GATK HaplotypeCaller command (McKenna et al., 2010). Next, we merged the mutations called across clones, creating a single file with all mutations across all genomes. We then filtered this list for high-confidence mutation calls, which screens out a large fraction of mutations. This resulted in a set of 1208 putative mutations. For each of these, we created screenshots of the aligned reads at the putative variant site using BamSnap (Kwon et al., 2021) and visually verified that a mutation was present. We required that the mutation be present in at least 3 reads and in over > 80% of aligned reads for haploids or > 40% for diploids. 732 of the putative mutations were verified. Mutations were annotated using SnpEff (Cingolani et al., 2012).

### Fitness Measurement

All pre-prepared pools and strains were removed from -80°C storage and grown overnight at 30°C in glucose-containing medium. The set of 480 sequenced clones was pooled with unevolved barcoded ancestors, 2-day glucose mutants and their ancestor, and 1-, 5-, and 1/5-day glucose mutants and their ancestor (see Table 1 for pool component details). This pool was then mixed at a 1:10 ratio with an unbarcoded neutral strain (GSY146 alpha) and this was used to inoculate cultures to begin the fitness measurement assays. Populations were propagated in triplicate in nine different conditions (27 total experiments), including Gly/Eth media with transfers regimes of 2, 4, 6, 8, and 10 days between transfers and glucose media with 1, 2, 3, and 5 days between transfers. All fitness measurement cultures were propagated for 5 growth cycles. Cells were cultured in 100 mL of Gly/Eth or glucose media at 30°C while shaking at 200 RPM in unbeveled flasks and a minimum of 400 µL were transferred between growth cycles. At the end of each growth cycle, cultures were monitored with an OD600 reading, Coulter Counter measurement, and plating cells (CFUs/mL) to track viability.

As in the evolution the transfer volume was adjusted for some of the longer transfer regimes where viability dropped, so that ∼5e7 cells were transferred. Following each growth cycle, 5 mL of leftover culture was removed, centrifuged, and resuspended in 2 mL of 17% glycerol. Five 400 µL aliquots were stored at -80°C. The remaining 95 mL of culture was divided into four 50 mL conical tubes, centrifuged, washed once with 2 mL of TE buffer, and then resuspended in 1 mL of 0.9M sorbitol.

### DNA extraction

DNA was extracted from each of the 135 (27 experiments x 5 timepoints) samples collected in the same way described above for the evolutions.

### PCR amplification of barcodes

To reduce bias and improve the consistency of amplification, we used an amended 1-step amplification procedure for the fitness measurement assays (Hays et al., 2023). We used a set of 12 reverse (ROS) and 10 forward (FOS) uniquely labeled primers to multiplex our samples for sequencing. PCR reactions were prepared on ice in 96-well PCR plates. Two reactions were prepared per sample as follows:

25 µL of Q5

2.5 µL 1 µM ROS

2.5 µL 1 µM FOS

0.75-1.75 µL of DNA

18.75-19.25 µL dH_2_0

PCR protocol:

1. 98°C 2 minutes

2. 98°C 10s

3. 60°C 20s

4. 72°C 15s

5. Repeat steps 2-4 20 times

6. Hold at 10°C

The volumes of water and DNA were adjusted so that approximately 500 ng of DNA were added to each reaction. The progress of the reactions was checked via Qubit measurement and additional cycles were added until the concentration of DNA read by the Qubit was at least 6 ng/µL. The duplicated samples were then mixed, and a small amount of each sample was run on a 2% agarose gel to ensure the barcode was amplified as desired. Finally, 4 samples of similar concentration were mixed and 50 µL of the mixed samples was run on a 2% agarose gel. The DNA was then extracted from the gel using the QIAquick Gel Extraction Kit (Qiagen, ID 28704) and uniquely labeled samples were mixed according to concentration so that each sample would be evenly represented in the mix. DNA was then sequenced on two NovaSeq sequencing lanes.

### Analyzing amplicon sequencing and chimera correction

Fastq files were demultiplexed by ROS-FOS primer combinations and the barcode sequence was extracted from each read. Each of the 1,834 unique barcodes included in the pool were counted in each sample using a perfect match approach. A barcode was able to be extracted from approximately 90% of reads.

To correct for index hopping on the flow cell (Kircher et al., 2012), which can result in barcodes with mislabeled primer indices, we estimated the number of index-hopped reads generated for each barcode in each sample by first estimating the overall index hopping rate. This was done by counting the reads that show up in primer index combinations that were not included in the prep and should have no counts. We counted all reads with those specific ROSz and FOSy indices and calculated the index hopping rate as:

Index hopping rate = count in samples that should be 0/(ROSz * (FOSy/total reads on lane))

This was done for each empty index combination on each sequencing lane and then averaged within each lane to calculate the index hopping rate specific to that sequencing run. Using this rate and the count of each BC, ROS and FOS index, and BC_ROS and BC_FOS association, we then calculated a corrected barcode count for each BC in each FOS-ROS combination using the following formula:

corrected_BCx_count = BCx_count – (ROSz_BCx_count * Index hopping rate * FOSy_count/total reads + FOSy_BCx_count * Index hopping rate * ROSz_count/total read) / 2

where:

BCx_count = count of BCx in sample FOSy_ROSz

FOSy_count = count of all reads with FOSy

ROSz_count = count of all reads with ROSz

FOSy_BCx_count = count of all reads with BCx and FOSy

ROSz_BCx_count = count of all reads with BCx and ROSz

We validated this correction by showing that we could predict the individual barcode counts in the samples that should be 0 using this method (Figure S15). We also compared the counts obtained from technical PCR replicates before and after correcting for index hopping, finding that the correlation between technical replicates was high and improved after correcting for index hopping (Figure S16).

### Fitness measurement analysis

The index hopping-corrected barcode counts from each sample were then arranged in time series for each of the 27 measurement experiments. Two under-sequenced timepoints were removed from the trajectories (glucose 1-day replicate 1 timepoint 2 and Gly/Eth 4-day replicate 2 timepoint 5). The barcode trajectories of the three unevolved ancestors were aggregated to improve their fitness estimates. Trajectories were then analyzed using FitSeq2 (Li et al. 2023). The Fitseq2 outputs are in fitness per cycle relative to the mean fitness of the measurement pool. We filtered the output of Fitseq2 to include only those fitness measurements with error less than 5 (in units of per-cycle fitness). We also sought to only include adaptive mutants in our analysis and removed all clones that had lower fitness than their ancestor.

### Quantification of performance in different growth phases

Next, for each replicate we calculated the performance metrics of interest (Table 3) for each clone and for aggregated ancestors. Error estimates were propagated from the underlying fitness measurements and no estimates with error greater than 5 (in units of per-cycle fitness) were included. Fitness change per hour was used as the measurement of lineage performance in the different growth phases. The mean performance was calculated across the 3 replicates and the error was propagated using the standard method:

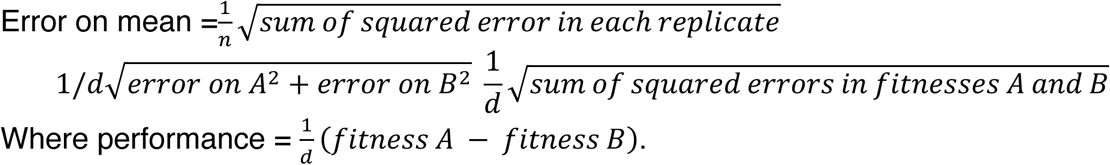

Note that *n* is the number of replicates begin averaged and *d* is the number of hours in the period during which the performance is being calculated.

The change in hourly performance in each part of the growth cycle was then calculated by subtracting the ancestral performance from the performance of evolved clones.

Change in mean performance metrics are then used in subsequent analyses.

## Data availability

All amplicon and whole genome sequencing data has been uploaded to the NIH Short Read Archive (SRA). The data has been assigned BioProject accession number PRJNA1394608.

Scripts and data for generating figures can be found at https://github.com/amahadevan99/stationary-phase-code.

## Acknowledgements

We thank the Stanford computing cluster for resources, and D. S. Fisher for useful discussions. We also thank the members of Sherlock lab for useful discussions, and particularly Katja Schwartz for help with troubleshooting of experimental protocols. This work was supported in part by grants from the NIH (NIGMS R35 GM131824) (G.S.), and the NSF (DMS-2235451) and Simons Foundation (MPS-NITMB-00005320) to the NSF-Simons National Institute for Theory and Mathematics in Biology (AM). This research was also supported in part by the National Science Foundation (NSF) via NSF PHY-2210386. AM acknowledges support from the Kavli Institute for Theoretical Physics (KITP) through the Gordon and Betty Moore Foundation Grants No. 2919.02, No. NSF PHY-1748958 and No. PHY-2309135, the Heising-Simons Foundation, and the Simons Foundation (216179, LB).

## Supplemental Figures

**Figure S1.**
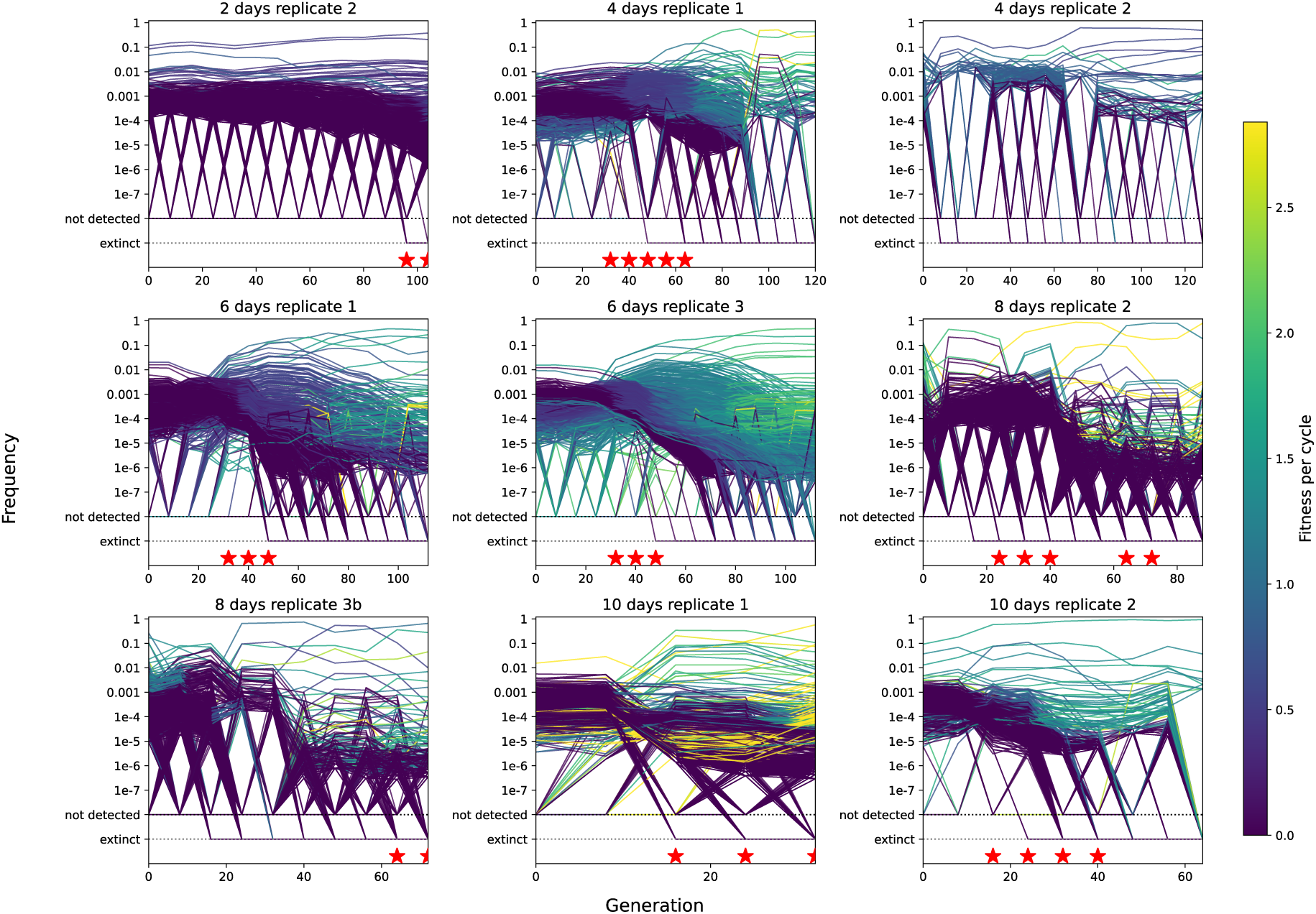
Evolutionary trajectories of a subset of Gly/Eth evolved populations. Each line shows the trajectory of a barcoded lineage colored based on the fitness inferred by FitMut2. Lineages are plotted as extinct when they are not detected in a timepoint and are also undetectable in all subsequent timepoints. Red stars indicate the timepoints from which clones were sampled.

**Figure S2.**
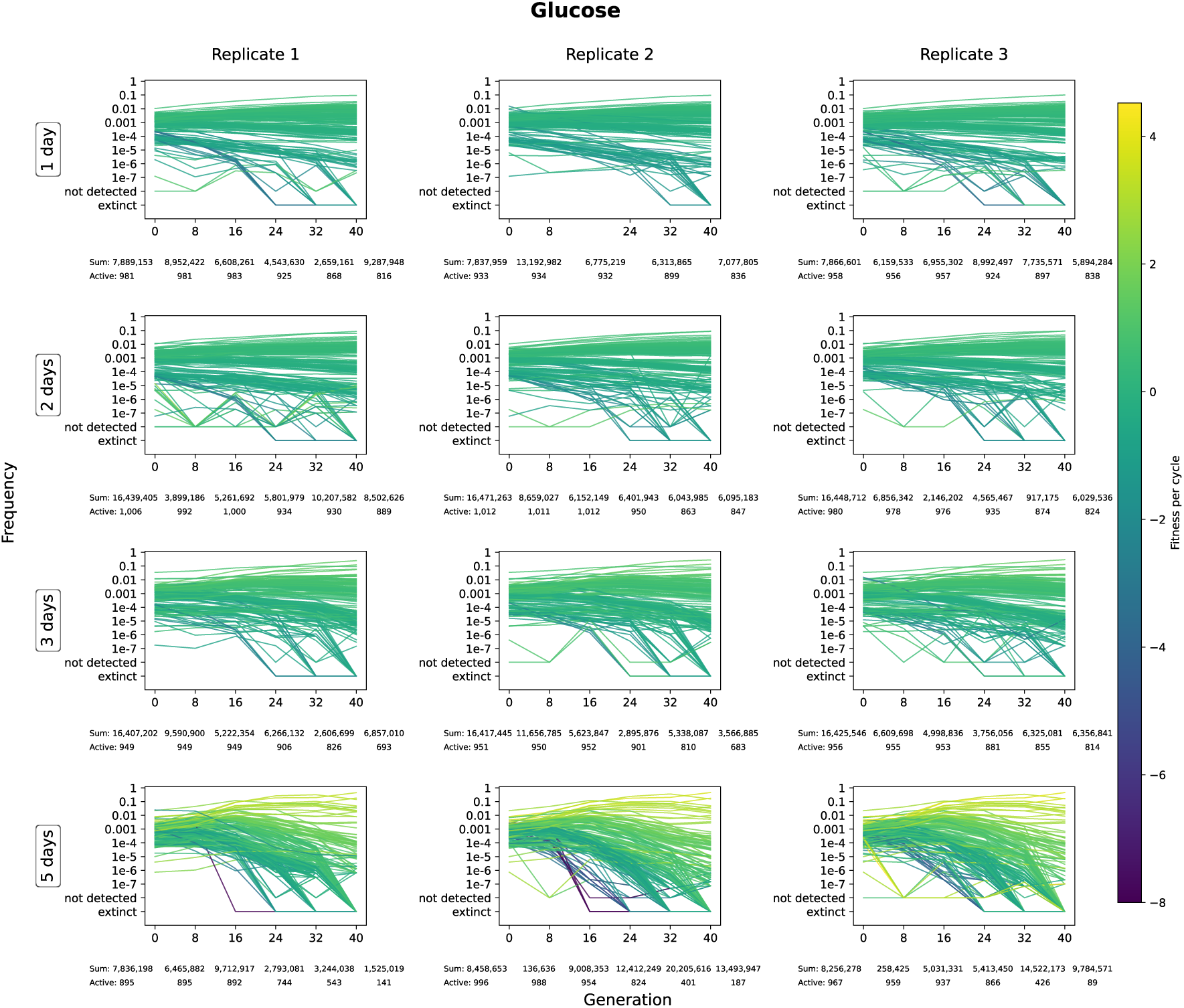
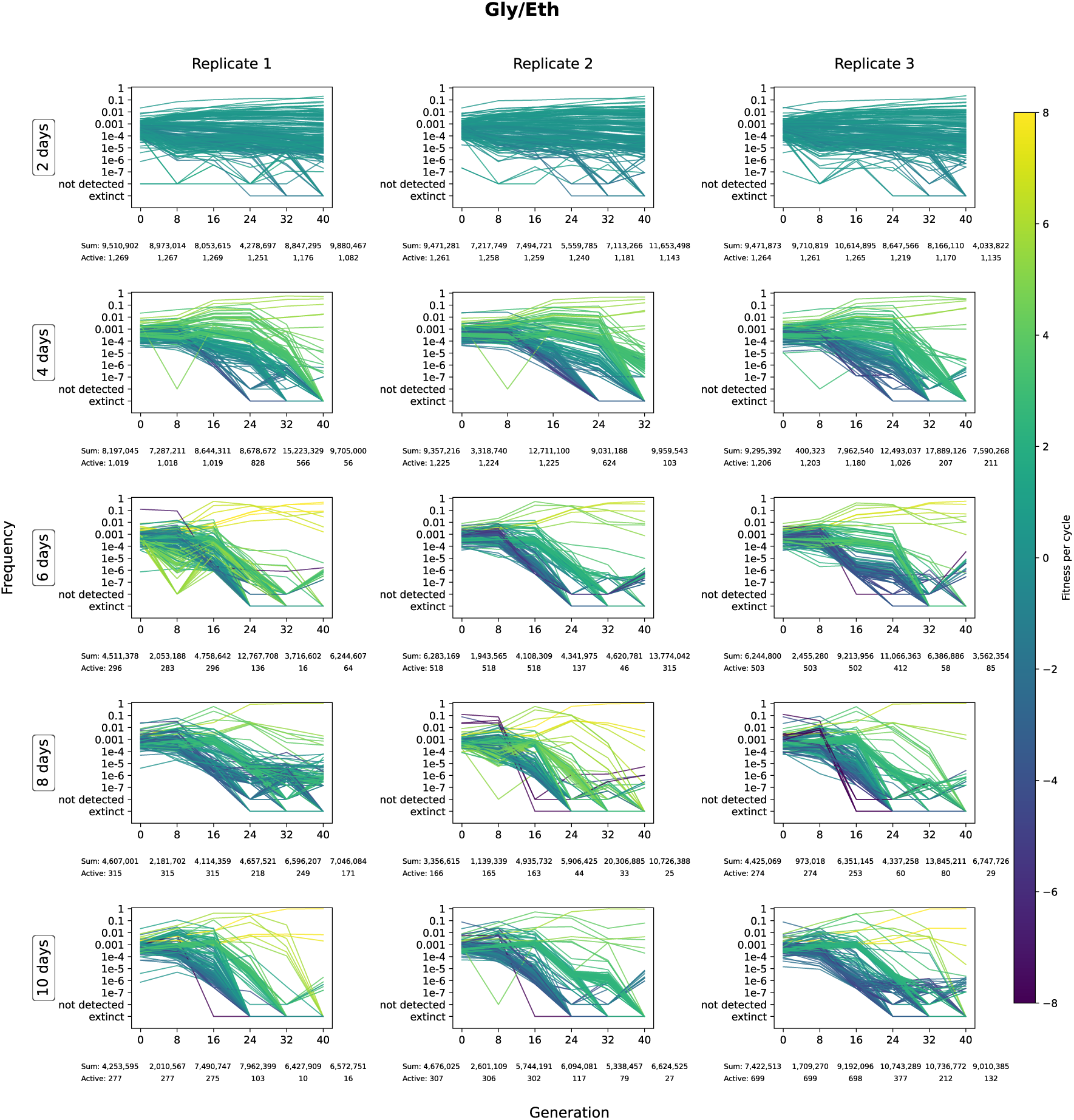
Fitness remeasurement trajectories. Trajectories of a subset of barcodes in the remeasurement pool are shown for all 3 replicates measured in 9 conditions. The top 100 fittest lineages and 100 randomly selected lineages are plotted for each replicate. Only lineages for which fitness is estimated with an error of <5 are plotted. Lineages are colored by the fitness inferred by FitSeq2. Sum indicates the total number of counts in each timepoint and Active indicates the number of non-zero unique barcodes in each timepoint.

**Figure S3.**
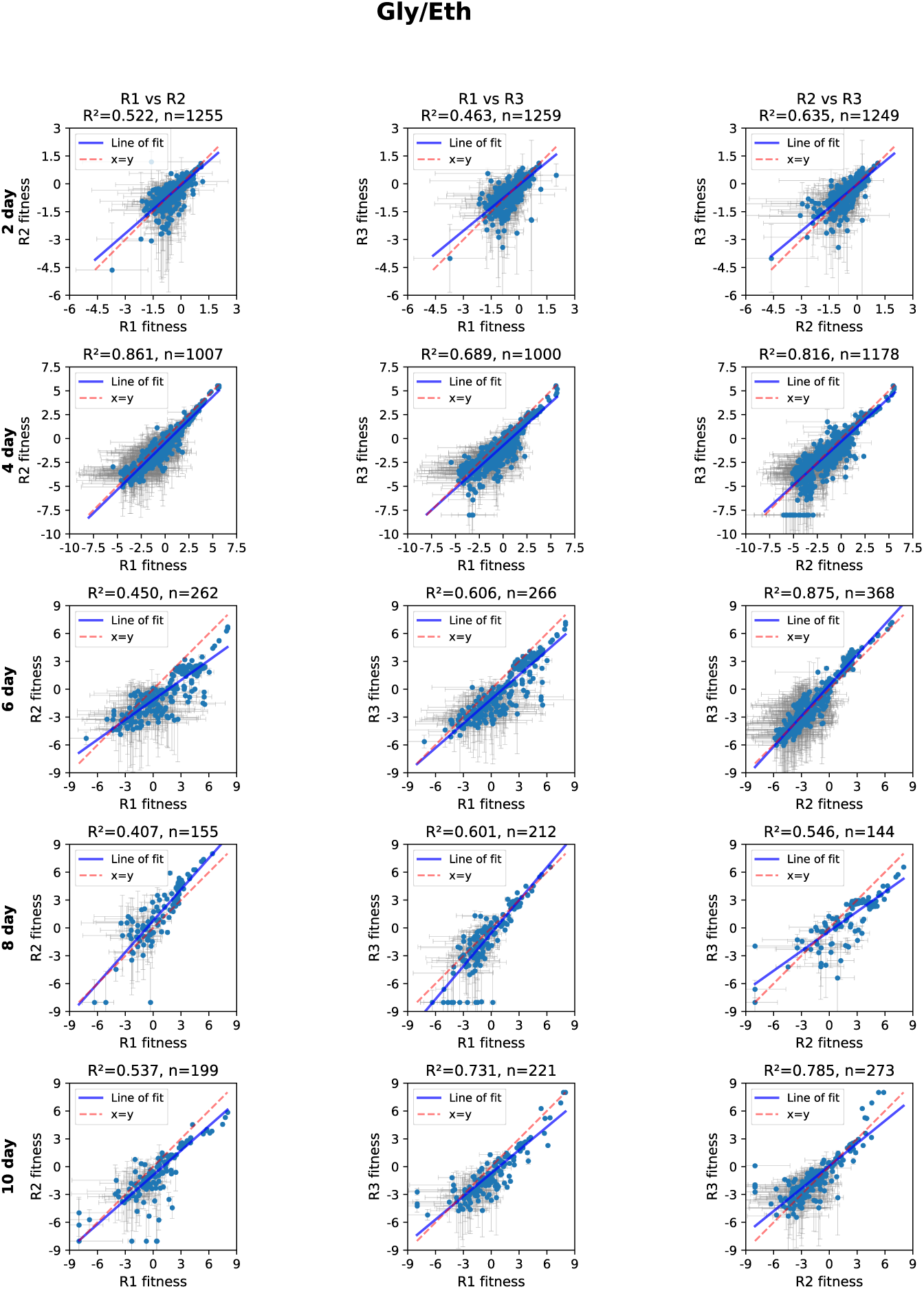
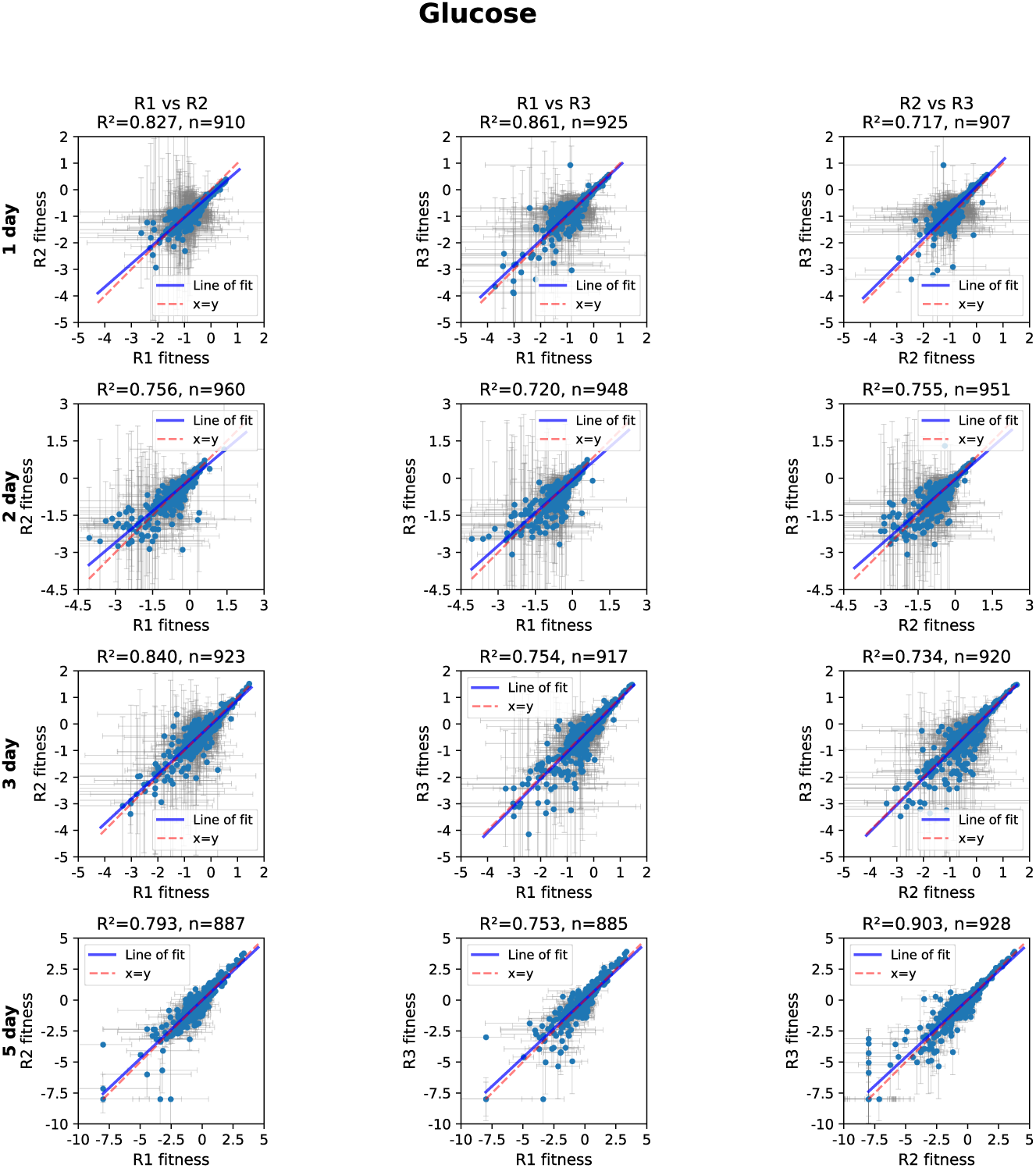
Replicate-replicate fitness correlations. For each assay condition replicate-replicate correlations are shown for fitness per cycle inferred by FitSeq2. Error bars shown in grey are estimated by FitSeq2. Only measurements with error less than 5 units are included in the plots and downstream analysis.

**Figure S4.**
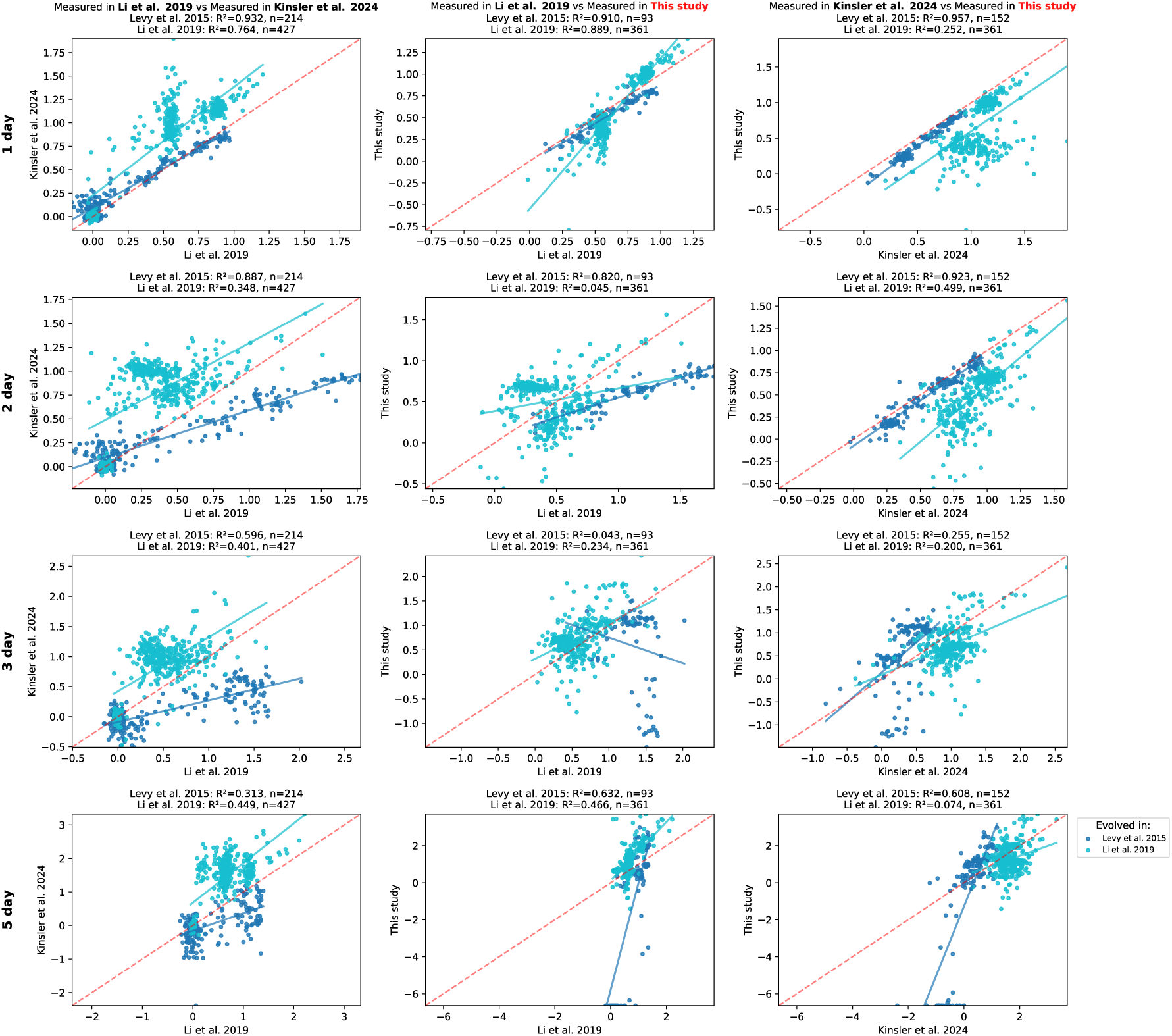
Measurements of glucose evolved clones in this study are comparable with previous measurements. Many of the glucose evolved clones included in our fitness remeasurement pool have been assayed previously in glucose conditions in Li et al. 2019 and Kinsler et al. 2024. The blue points show 2 day evolved glucose clones (Levy et al. 2015) and the cyan points show 1, 5, and 1/5 day glucose evolved clones (Li et al. 2019).

**Table S1.** Ras/PKA mutations are less common in evolutions that include a stationary phase. The number of mutations in genes in the Ras/PKA pathway in clones isolated from different evolutionary conditions is shown.

| Gene | Gly/Eth |  |  |  |  | 5 day<br>Gluc. |
| --- | --- | --- | --- | --- | --- | --- |
|  | 2 day | 4 day | 6 day | 8 day | 10 day |  |
| *IRA1* | 15 | 1 | 2 | 0 | 0 | 0 |
| *IRA2* | 17 | 1 | 3 | 0 | 2 | 1 |
| *PDE1* | 0 | 0 | 0 | 0 | 0 | 0 |
| *PDE2* | 8 | 0 | 0 | 0 | 0 | 0 |
| *GPB1* | 1 | 0 | 0 | 0 | 0 | 0 |
| *GPB2* | 8 | 0 | 0 | 1 | 2 | 0 |
| *YAK1* | 7 | 3 | 1 | 0 | 1 | 0 |
| *RAS1* | 0 | 0 | 0 | 0 | 0 | 1 |
| *RAS2* | 0 | 0 | 0 | 0 | 1 | 0 |
| *CYR1* | 1 | 0 | 0 | 0 | 0 | 1 |
| *TPK1* | 1 | 0 | 0 | 0 | 0 | 0 |
| *MSN4* | 1 | 0 | 0 | 0 | 0 | 0 |
| *BCY1* | 1 | 0 | 0 | 0 | 0 | 0 |
| *GPA2* | 0 | 0 | 0 | 0 | 1 | 0 |
| Clones with<br>sufficient<br>coverage in WGS<br>(see Methods) | 86 | 81 | 179 | 49 | 54 | 71 |

**Figure S5.**
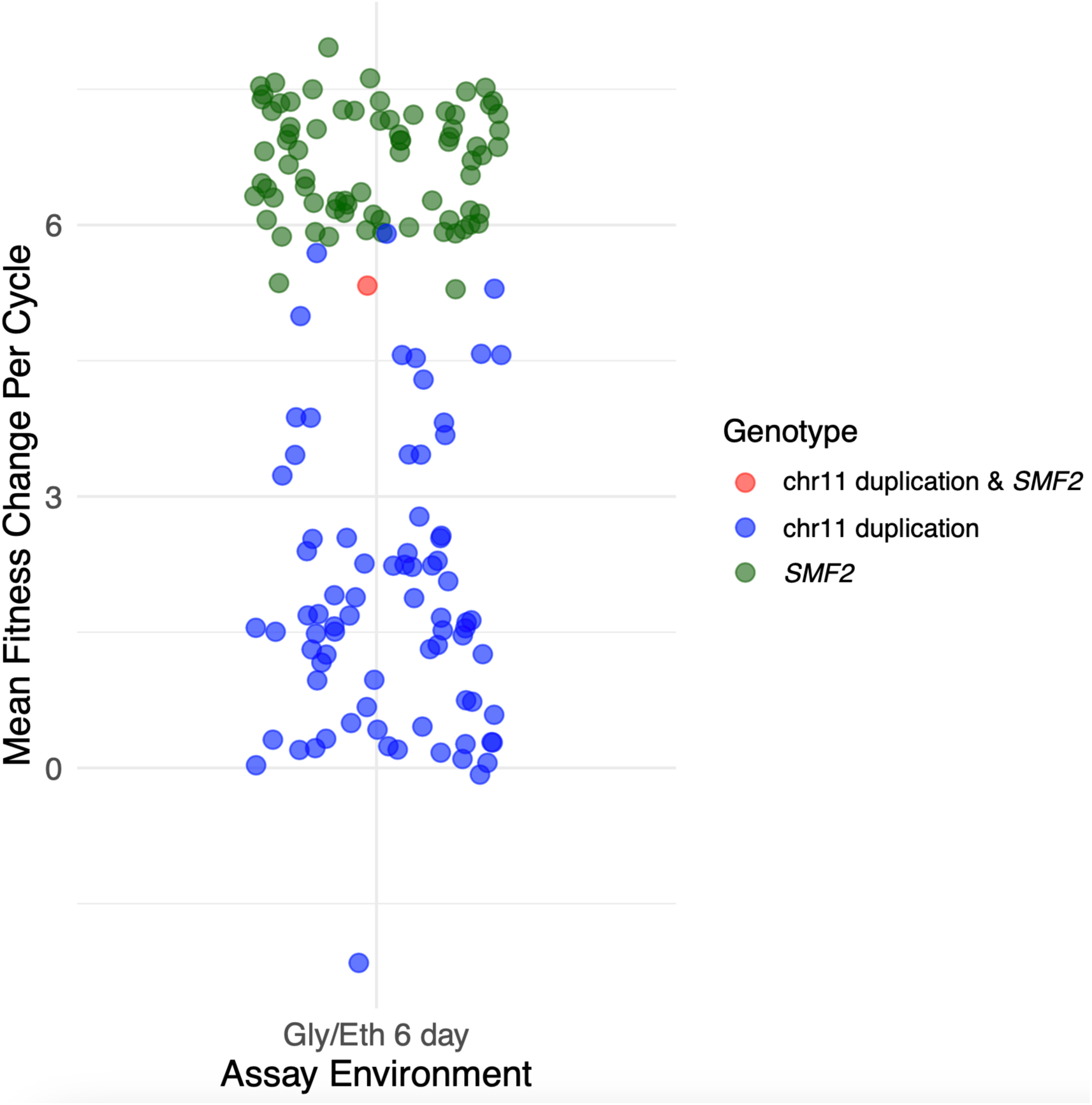
The *SMF2* mutants with a chromosome 11 duplication have lower fitness than other *SMF2* mutants. The Gly/Eth 6 day fitness of all evolved clones with a chromosome 11 duplication and/or an *SMF2* mutation. The chromosome 11 duplication does not provide any additional benefit in the *SMF2* mutant background.

**Figure S6.**
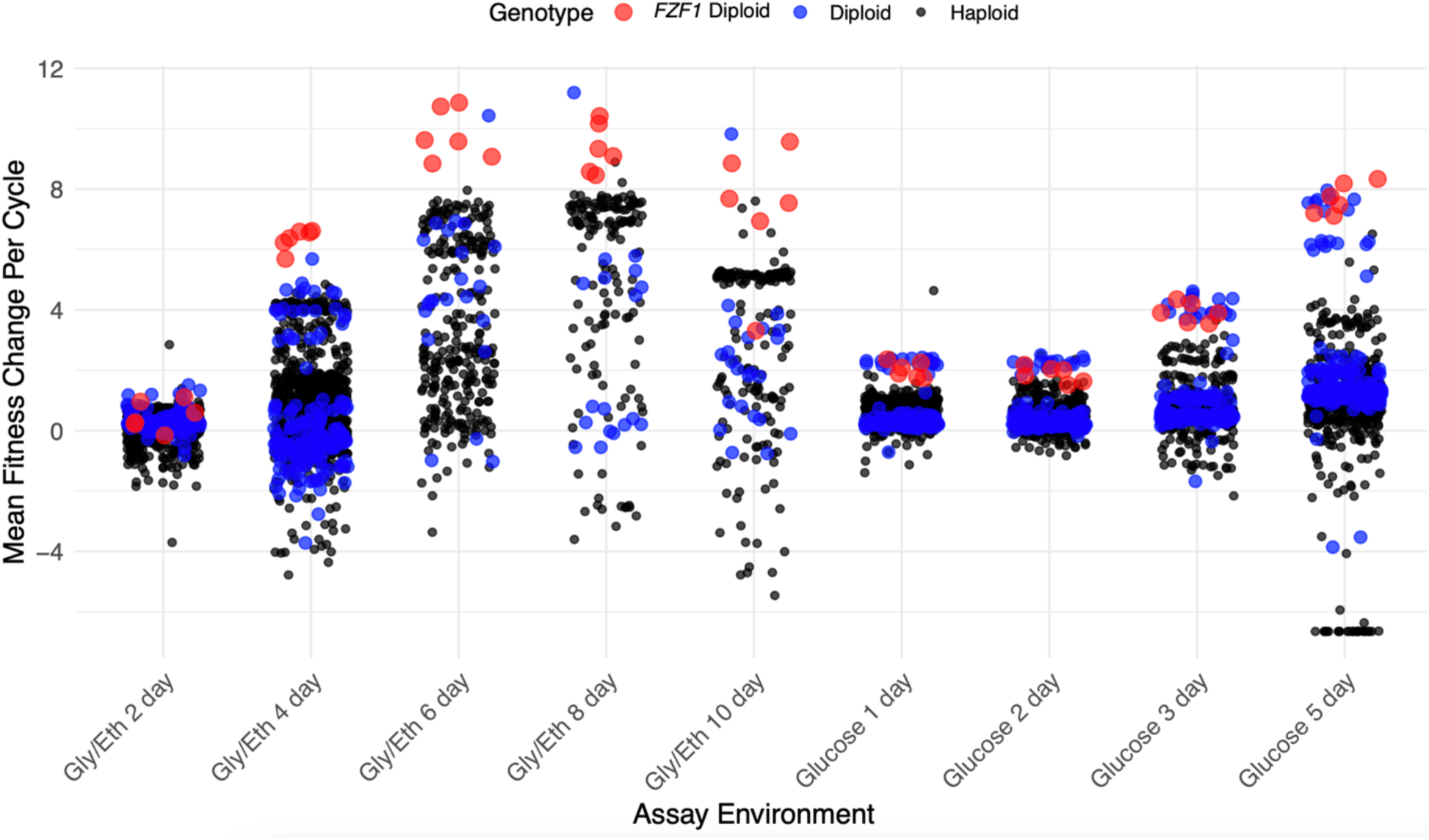
*FZF1* mutations are among the fittest in all conditions with a stationary phase. The mean fitness change per cycle of all adaptive mutants is plotted for each of the remeasurement conditions. The *FZF1* mutants, all of which are diploid, are colored red and perform particularly well in Gly/Eth conditions with stationary phase. Note that one of the *FZF1* clones was poorly measured and is not included in the plot.

**Figure S7.**
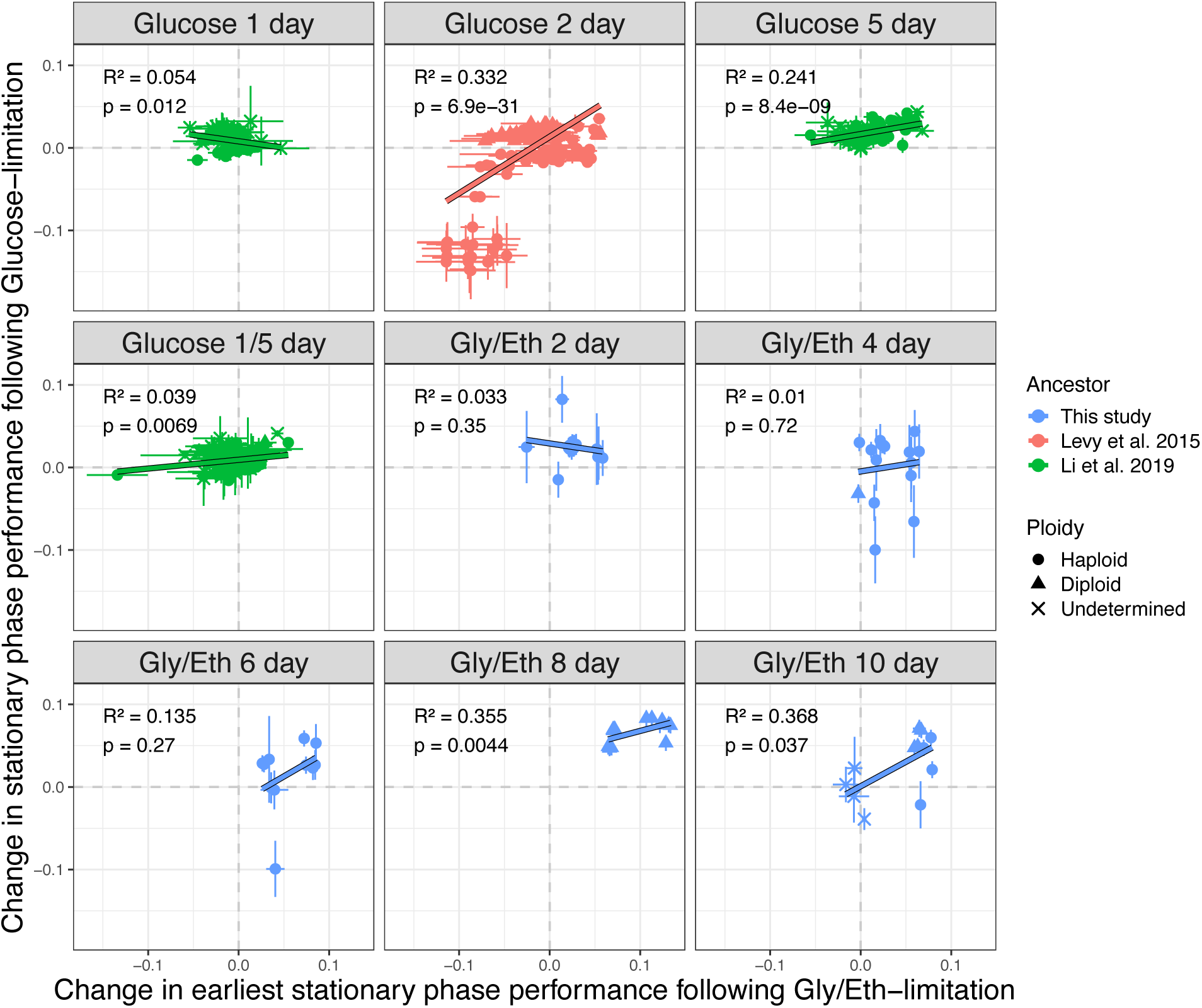
Stationary phase performance is correlated following glucose and Gly/Eth-limitation for clones that evolved with a stationary phase. Change in stationary phase performance (day 3-5) following glucose-limitation are correlated with changes in stationary phase performance (day 2-4) following Gly/Eth-limitation. For each evolution condition that included stationary phase, we observed a positive correlation between stationary phase performance following glucose and Gly/Eth-limitation.

**Figure S8.**
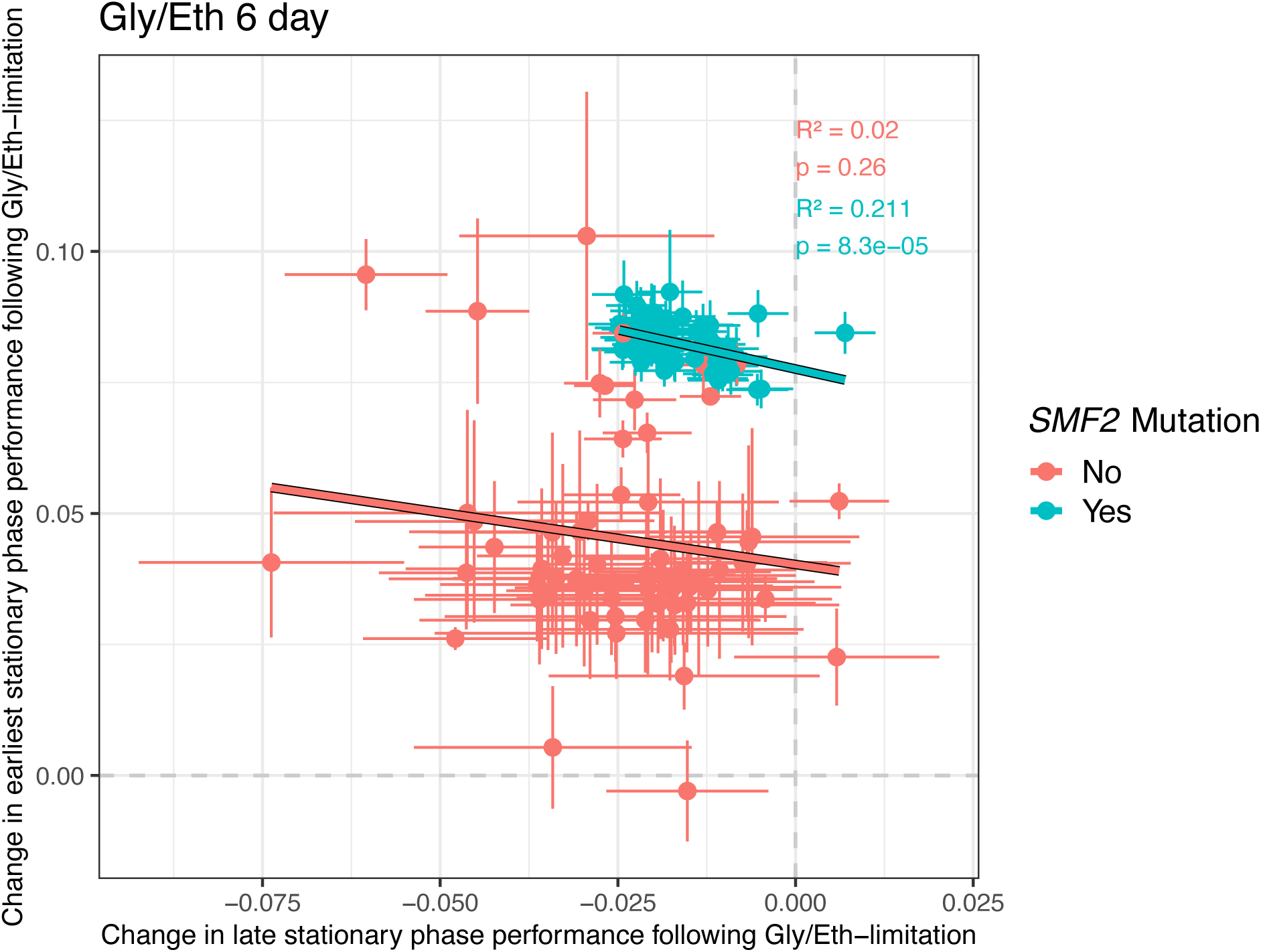
*SMF2* mutants cluster in trait-space compared to other adaptive mutants. The change in earliest (day 2-4) and late (day 6-10) stationary phase following Gly/Eth-limitation is shown for Gly/Eth 6-day adaptive haploid mutants. Clones in which an *SMF2* mutation was detected are colored turquoise and all other adaptive clones are shown in red. Results of a Pearson correlation test are shown for each group of adaptive mutants. Error bars indicate the combined error inferred by Fitseq2 for each underlying fitness measurement.

**Figure S9.**
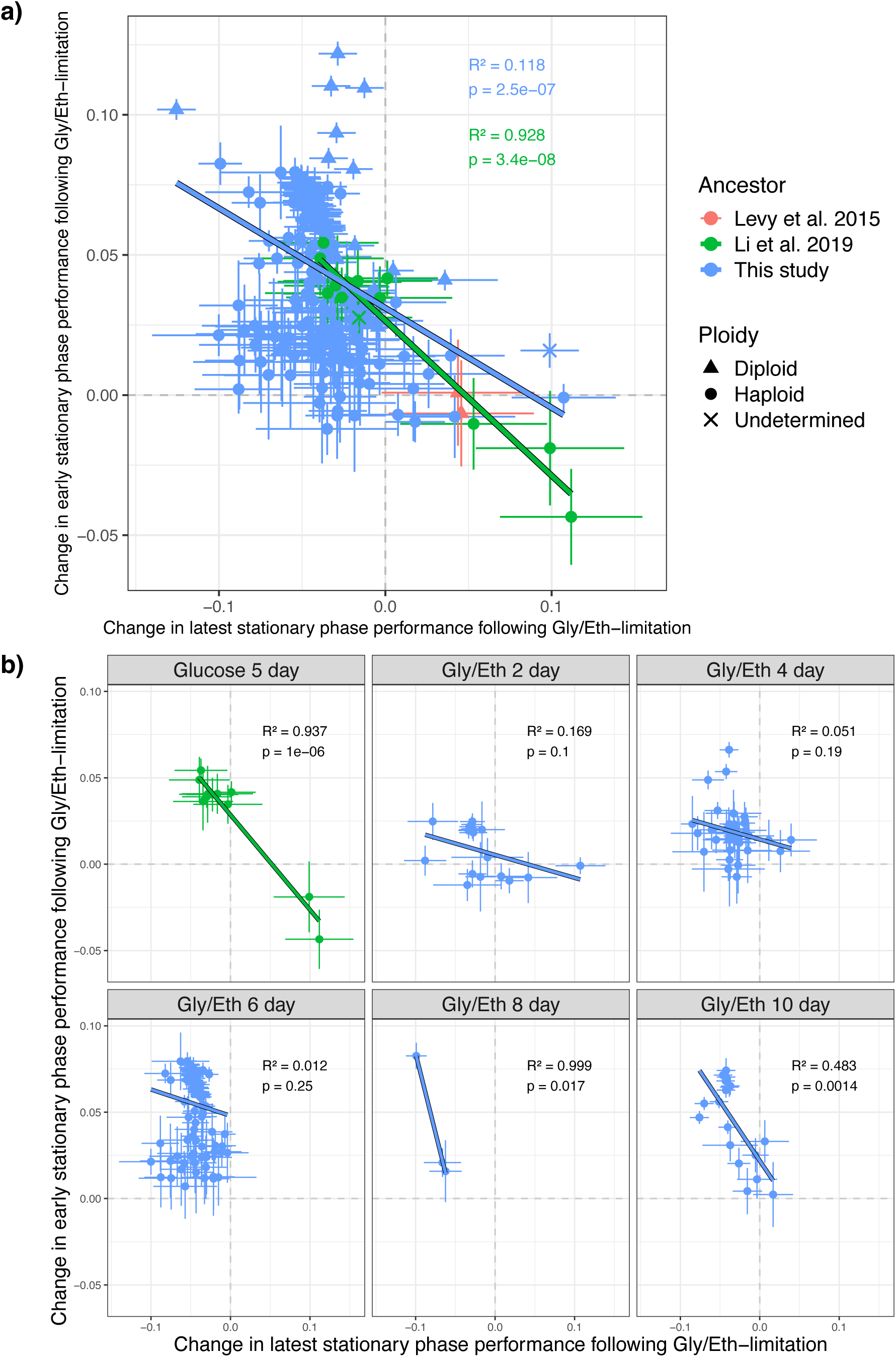
A trade-off between early and latest stationary phase performance. Adaptive haploids are plotted. A linear regression is shown through each set of all adaptive mutants comparing changes in <u>early</u> stationary phase performance (day 2-6) to <u>latest</u> stationary phase performance (day 8-10). (b) A linear regression is shown for adaptive haploid clones isolated from each evolution condition. Points are colored by the ancestor. Results of a Pearson correlation test are shown for each evolved group of adaptive haploid mutants. Error bars indicate the combined error inferred by Fitseq2 for each underlying fitness measurement.

**Figure S10.**
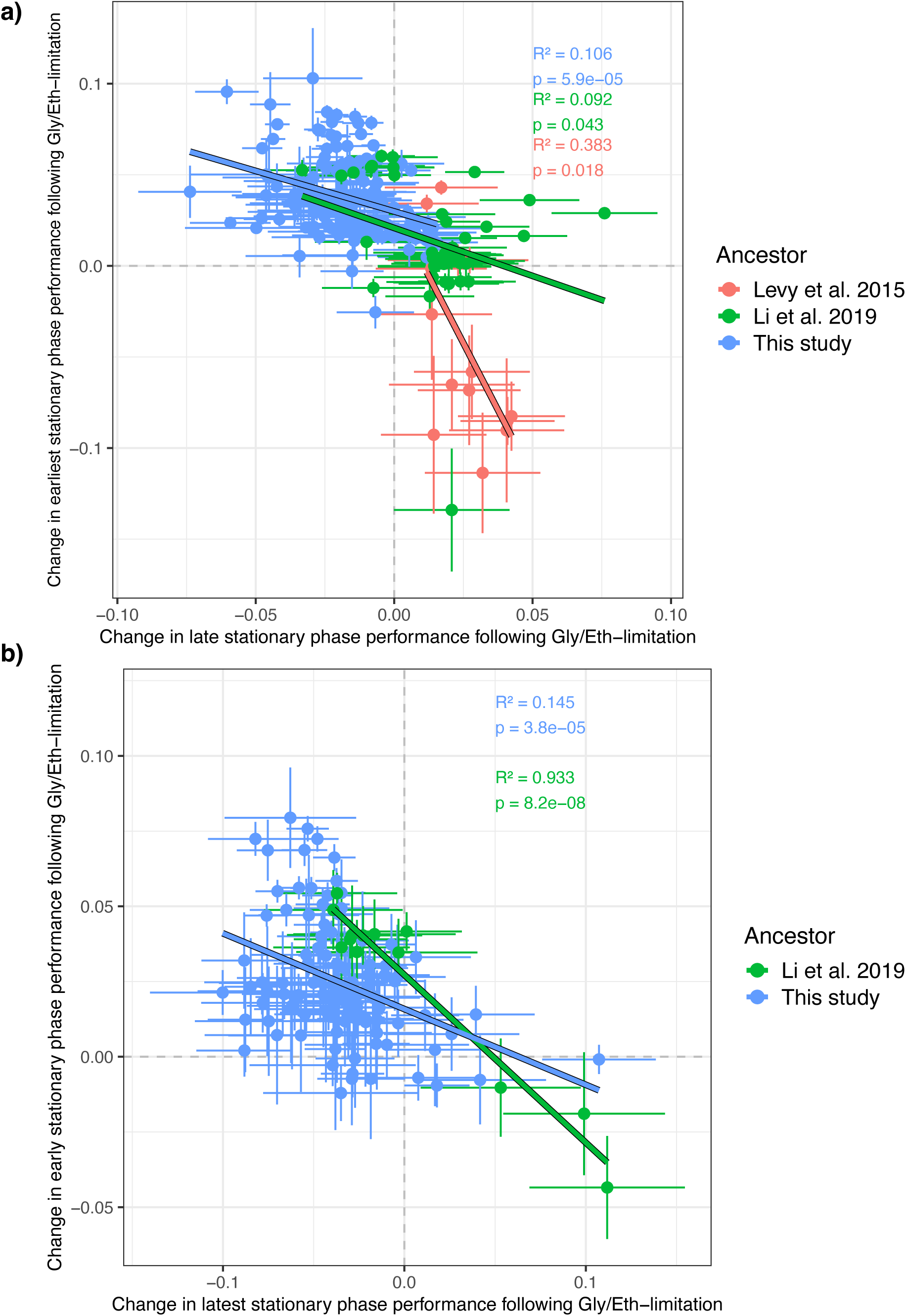
A trade-off between late and early stationary phase performance for adaptive haploids without *SMF2* mutations. (a-b) A linear regression is shown through each set of adaptive haploid mutants that lack the *SMF2* mutation comparing changes in <u>early</u> stationary phase performance (day 2-6) to <u>latest</u> stationary phase performance (day 8-10) (a) and comparing <u>earliest</u> stationary phase performance (day 2-4) to <u>late</u> stationary phase performance (day 6-10) (b). Results of a Pearson correlation test are shown for each group of adaptive mutants. Error bars indicate the combined error inferred by Fitseq2 for each underlying fitness measurement.

**Figure S11.**
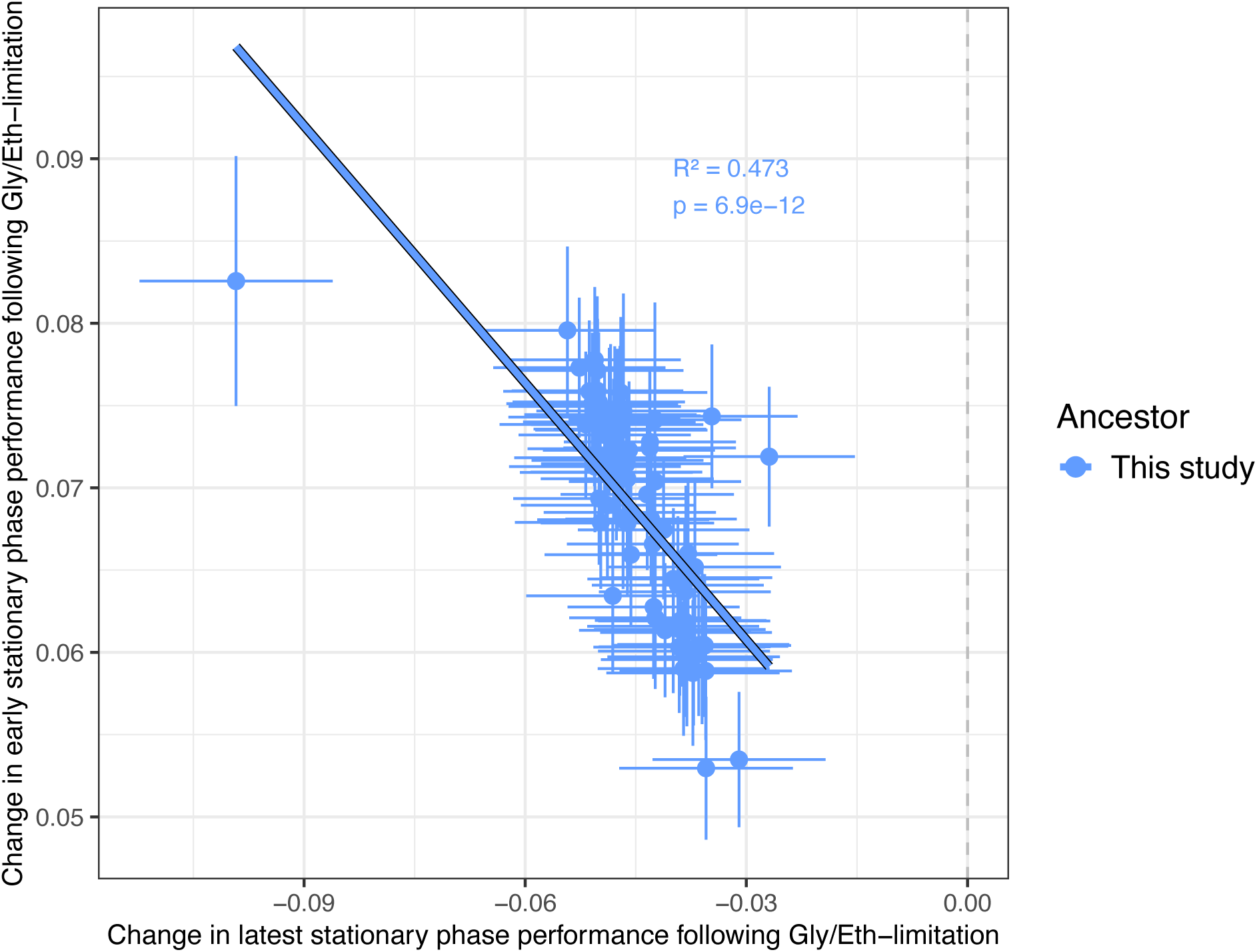
Trade-offs between early and late stationary phase performance among *SMF2* mutants. Changes in <u>early</u> stationary phase performance (day 2-6) following Gly/Eth-limitation are negatively correlated with changes in <u>latest</u> stationary phase (day 8-10) following Gly/Eth limitation. Only adaptive haploid mutants with a normal karyotype and an *SMF2* mutation are plotted. Results of a Pearson correlation test are shown.

**Figure S12.**
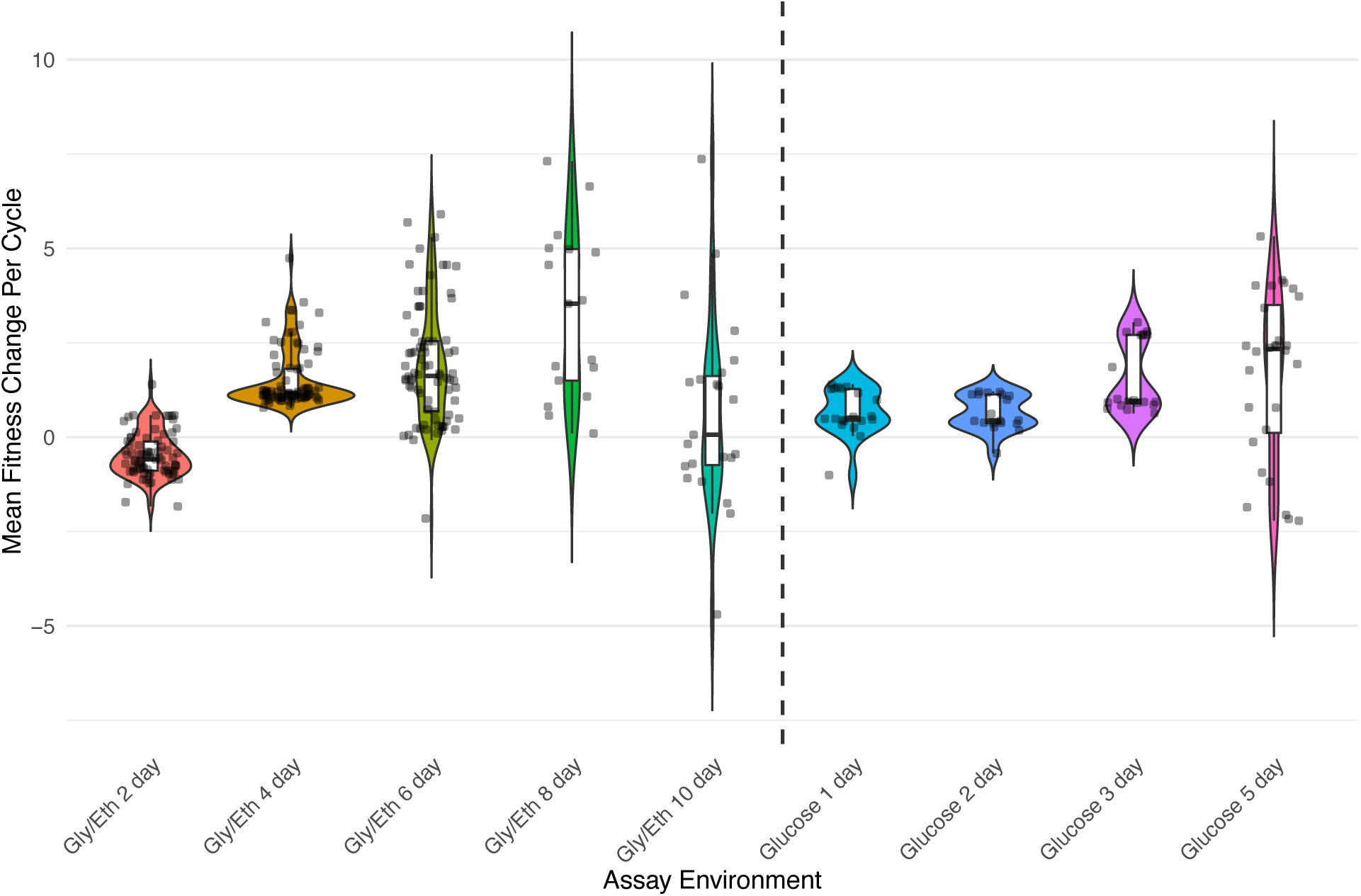
Most adaptive mutants with chromosome 11 duplications have increased fitness in the Gly/Eth 6 day condition. The change in mean fitness per cycle is plotted for all clones with a chromosome 11 duplication. These duplications are often deleterious in the 2-day and 10-day condition. There are fewer points plotted in some conditions (e.g. Gly/Eth 8 day and 10 day) because the clones drop in frequency in the fitness remeasurement assays resulting in the FitSeq2 fitness estimates having large error and not passing our filter.

**Figure S13.**
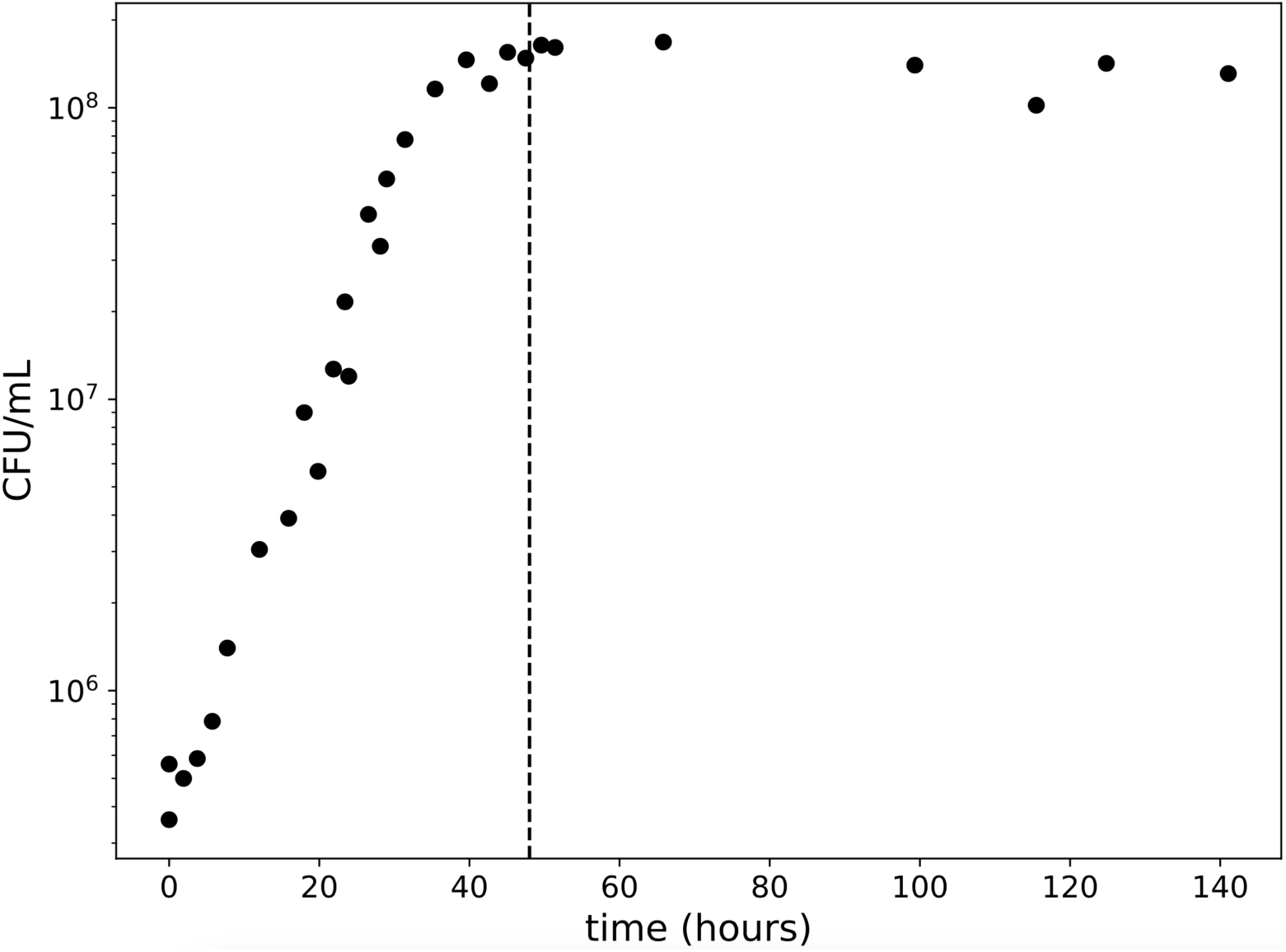
Barcoded yeast library reaches stationary phase after 48 hours. The barcoded yeast library was inoculated into 100 mL of Gly/Eth media and CFUs were measured by plating every ∼2 hours. The dashed line shows the 48-hour timepoint after which the number of viable cells stops increasing.

**Figure S14.**
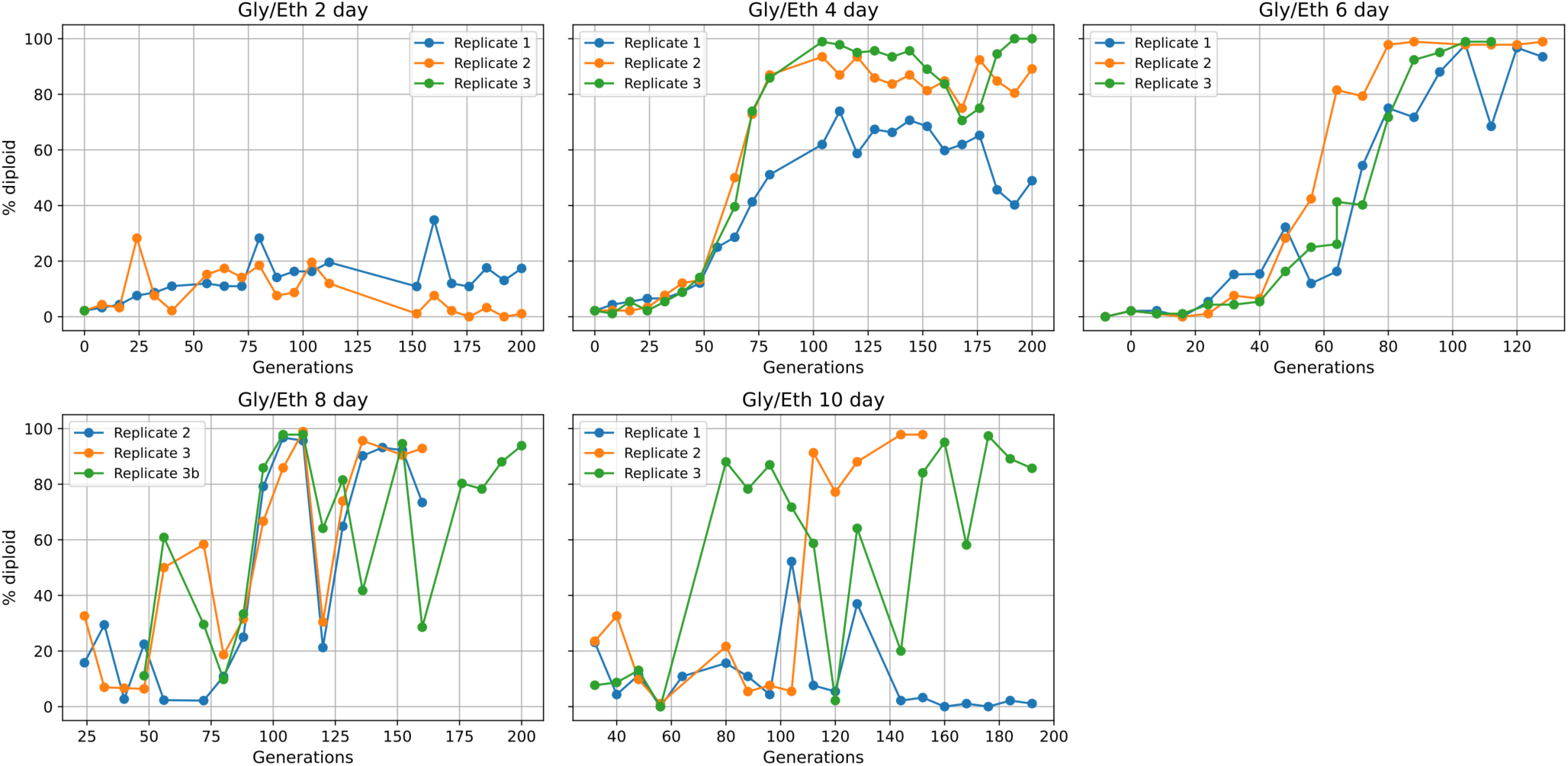
Diploids are selected during evolution in Gly/Eth. We measured the percent of cells that were diploid using benomyl assays as populations evolved. Each panel shows replicate populations from different evolution conditions.

**Figure S15.**
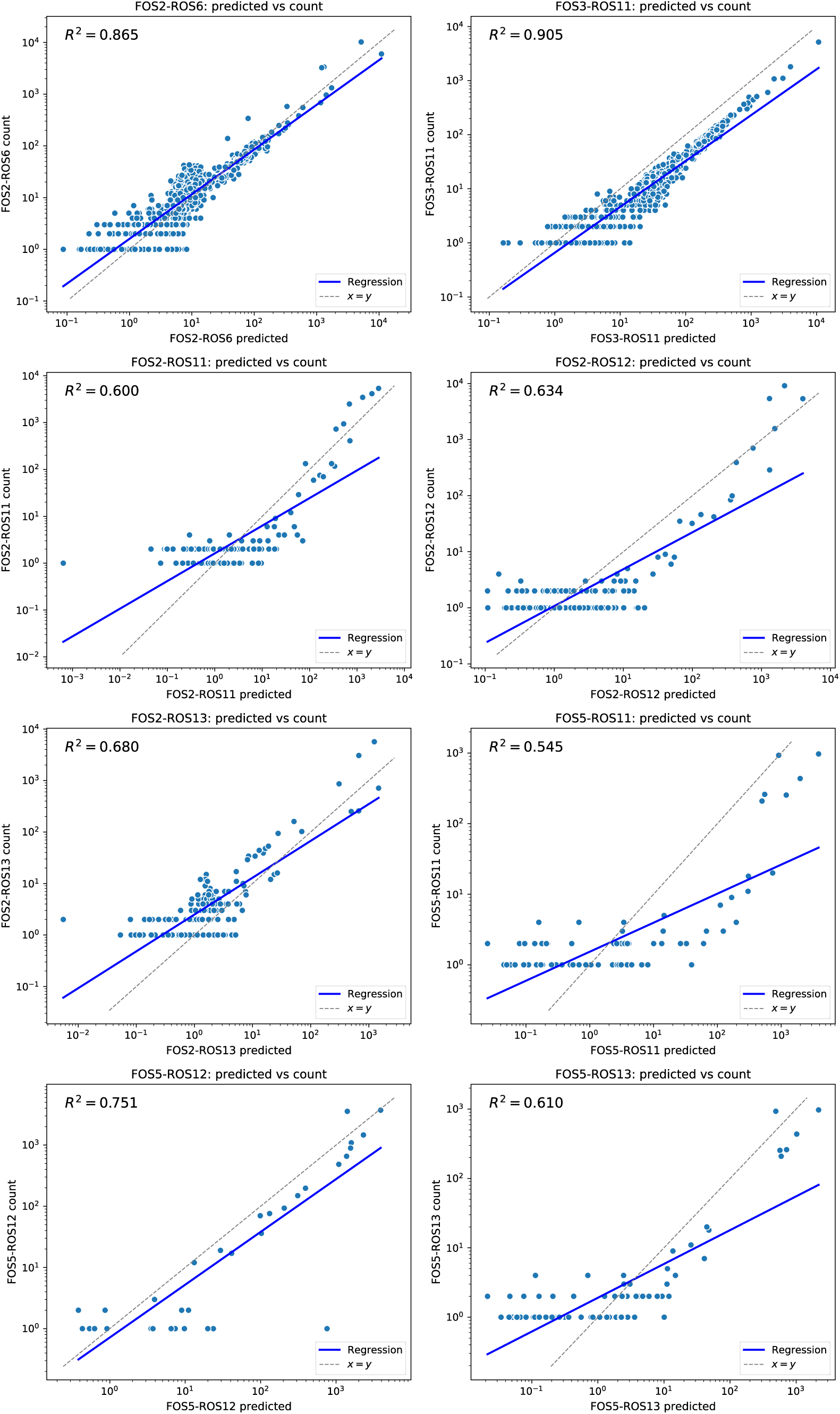
Generation of index hopped reads can be predicted. Eight FOS-ROS combinations were not included in the library preparation. These barcode counts are shown on the y-axis of each plot. The predicted number of BC for that FOS-ROS combination is shown on the x-axis. We corrected all counts to account for the generation of index hopped reads (see Methods).

**Figure S16.**
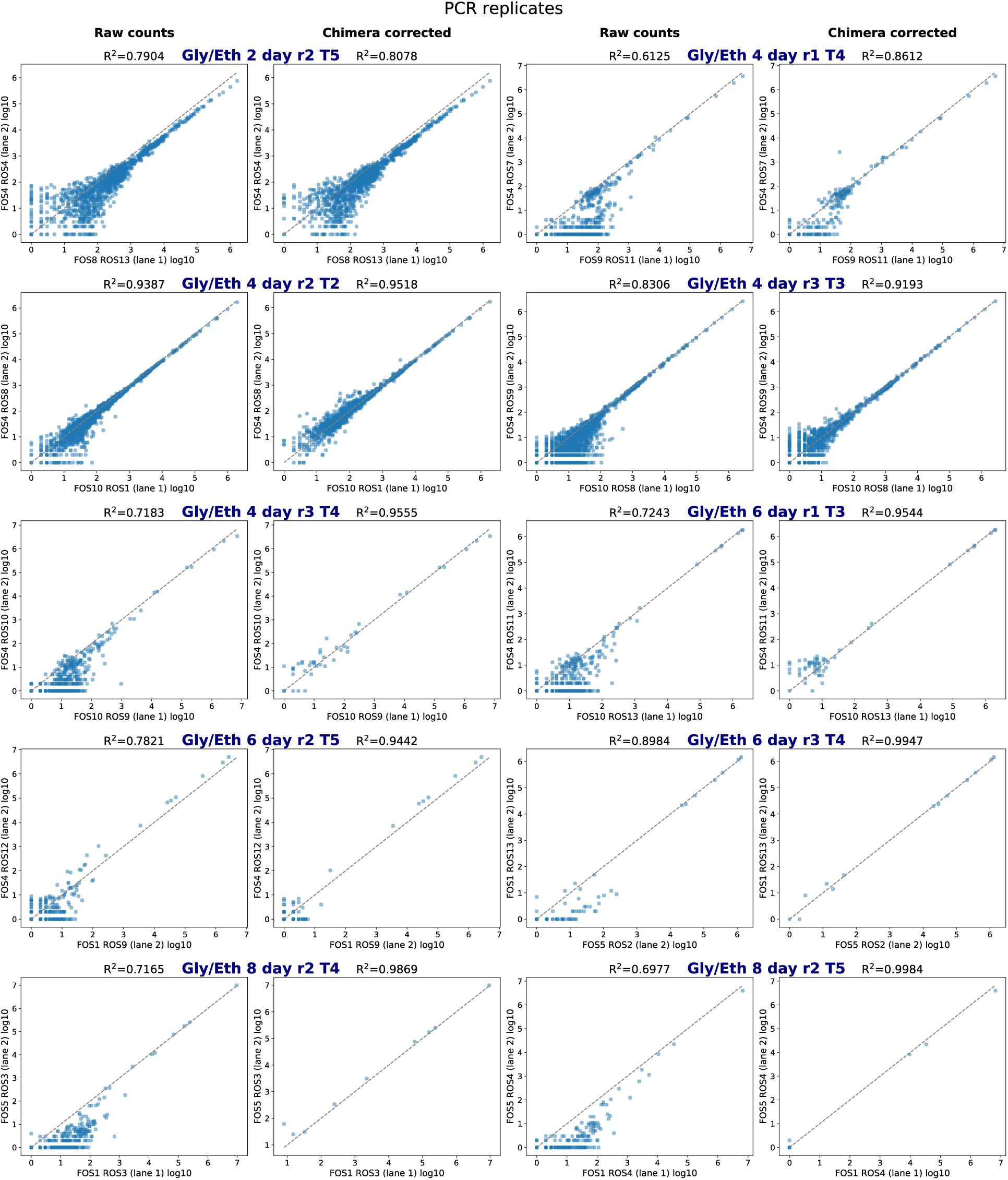
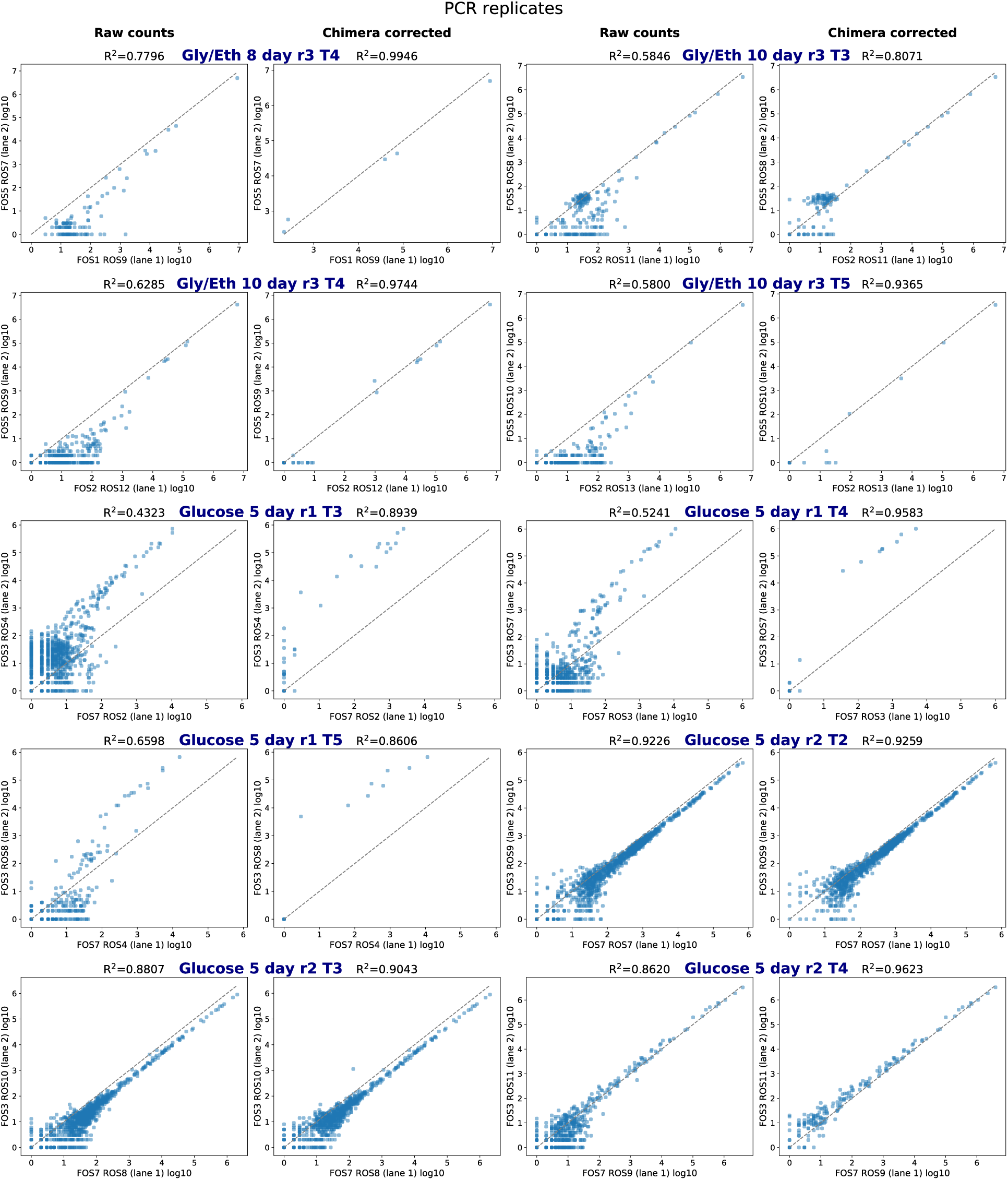
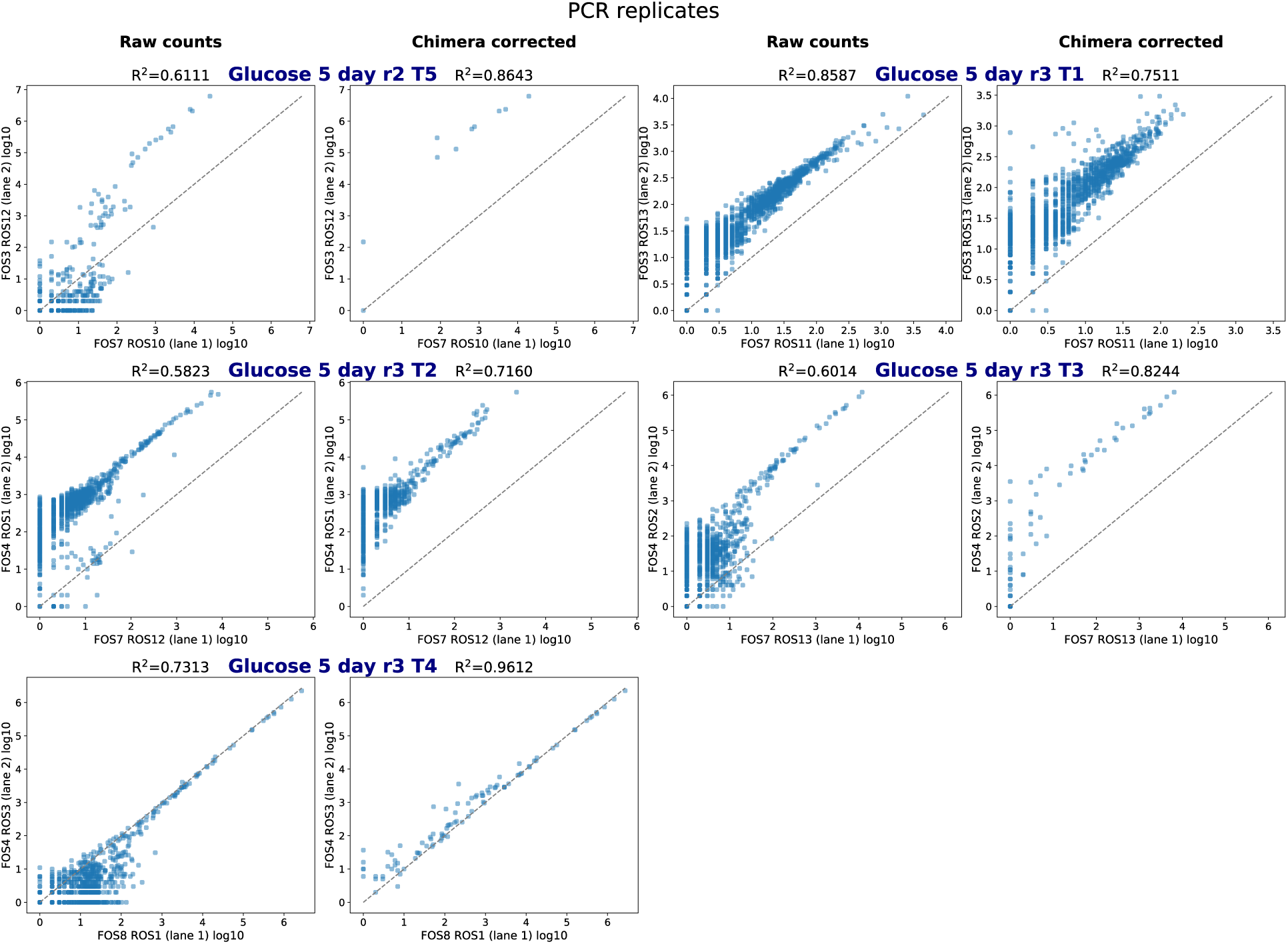
Technical PCR replicates are correlated. For a handful of samples we prepared a second amplicon library for quality assessment purposes. Here we show the raw and index-hopped correct counts from each of the library preparations (technical PCR replicates). Following correction for index-hopping, technical replicates are highly correlated even when one of the amplicon libraries is under-sequenced.

